# Genomic analyses of the *Linum* distyly supergene reveal convergent evolution at the molecular level

**DOI:** 10.1101/2022.05.27.493681

**Authors:** Juanita Gutiérrez-Valencia, Marco Fracassetti, Emma L. Berdan, Ignas Bunikis, Lucile Soler, Jacques Dainat, Verena E. Kutschera, Aleksandra Losvik, Aurélie Désamoré, P. William Hughes, Alireza Foroozani, Benjamin Laenen, Edouard Pesquet, Mohamed Abdelaziz, Olga Vinnere Pettersson, Björn Nystedt, Adrian Brennan, Juan Arroyo, Tanja Slotte

## Abstract

Supergenes govern balanced polymorphisms in a wide range of systems. The reciprocal placement of stigmas and anthers in pin and thrum floral morphs of distylous species constitutes an iconic example of a balanced polymorphism governed by a supergene, the distyly *S-*locus. Recent studies have shown that the *Primula* and *Turnera* distyly supergenes are both hemizygous in thrums, but it remains unknown if hemizygosity is pervasive among distyly *S*-loci. Here we have characterized the genetic architecture and evolution of the distyly supergene in *Linum* by generating a chromosome-level genome assembly of *Linum tenue*, followed by the identification of the *S*-locus using population genomic data. We show that hemizygosity and thrum-specific expression of *S*-linked genes, including a pistil-expressed candidate gene for style length, are major features of the *Linum S*-locus. Structural variation is likely instrumental for recombination suppression, and although the non-recombining dominant haplotype has accumulated transposable elements, *S-*linked genes are not under relaxed purifying selection. Our findings reveal remarkable convergence in the genetic architecture and evolution of independently derived distyly supergenes. The chromosome-level genome assembly and detailed characterization of the distyly *S*-locus in *L. tenue* will facilitate elucidation of molecular mechanisms underlying the different forms of flowers described by Darwin.

## Introduction

Supergenes control complex phenotypic polymorphisms under balancing selection through the preservation of advantageous allelic combinations (Thompson and Jiggins 2014). They occur in a wide range of systems, including ants, butterflies, and plants (reviewed in Schwander et al. 2014; Gutiérrez-Valencia et al. 2021). Distyly in flowering plants is one of the most emblematic instances of a multi-trait polymorphism controlled by a supergene termed the *S*-locus. Both the adaptive significance and inheritance of distyly (Bateson and Gregory 1905; Ernst 1936) have received sustained interest ever since Darwin’s studies (Darwin 1863; 1877).

In distylous lineages, individuals exhibit one of two types of flowers which differ primarily in the positions of their sexual organs (Fig. 1). While pin (L-morph) individuals are long-styled and present low anthers, thrum (S-morph) individuals are short-styled with anthers at a high level in the flower. Distyly has been suggested to increase the efficacy of pollination (Darwin 1877; Barrett 2002; Keller et al. 2014) through the deposition of pollen grains from each morph on different regions of pollinating insects’ bodies (Darwin 1877). Distyly has evolved convergently multiple times (Ganders 1979; Lloyd and Webb 1992), and is frequently associated with a heteromorphic self-incompatibility system (Dulberger 1992; Barrett 2002; 2019), which prevents inbreeding and guarantees disassortative mating.

**Fig 1.**
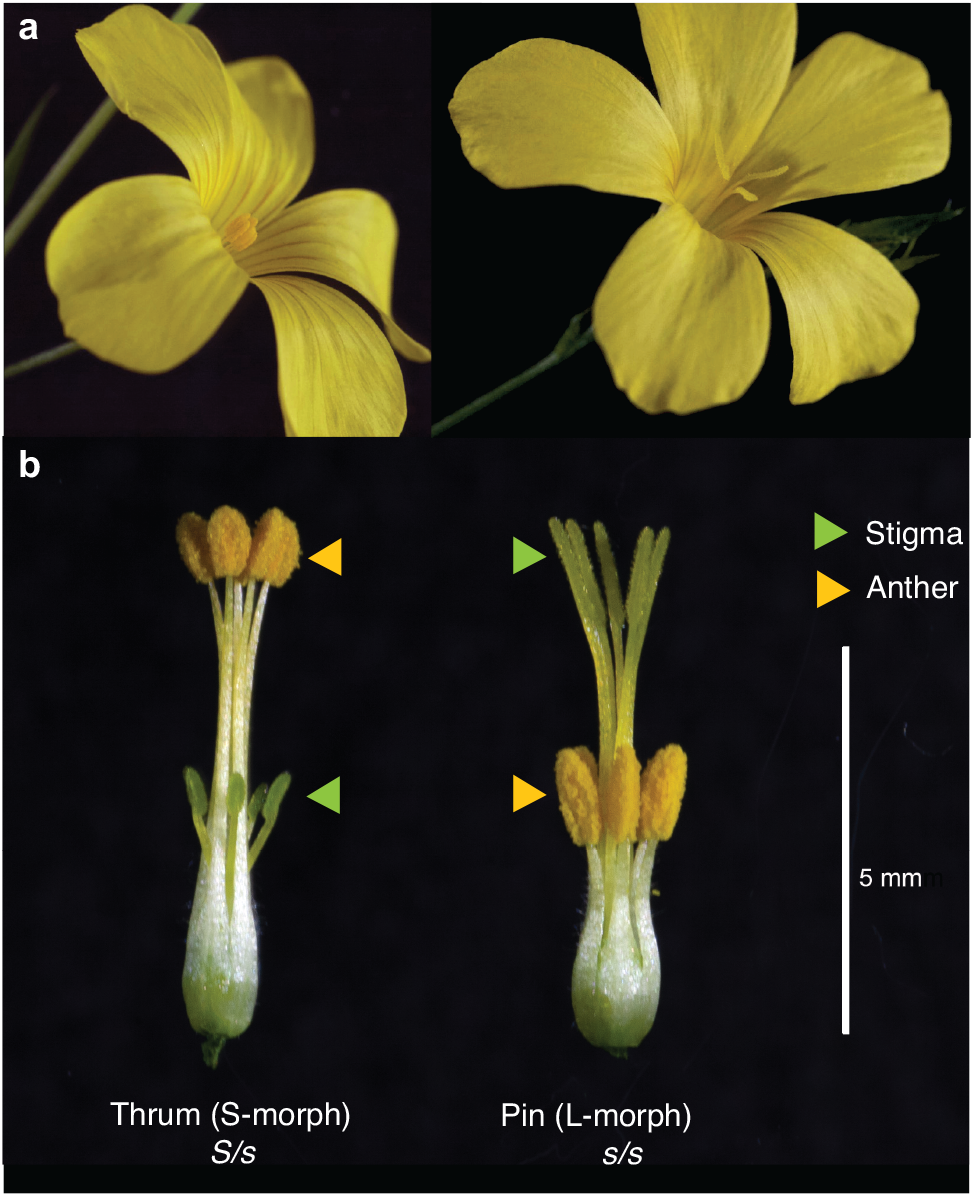
Distyly in *Linum tenue*. **a**, Thrum individuals (also termed S-morph, to the left) are short-styled while pin individuals are long-styled (L-morph, to the right). **b**. Reciprocal location of male (anther) and female (stigma) reproductive organs in thrum (left) and pin (right) flowers. Petals and sepals have been removed to better visualize reproductive structures. The expected *S-*locus genotype is indicated below each morph.

Early geneticists showed that distyly was inherited as if it was governed by a single Mendelian locus, with a short style allele that was dominant over the long style allele in several distylous species (Bateson and Gregory 1905; Laibach 1923; reviewed in Ganders 1979). The classic model of the *Primula* distyly *S*-locus (Ernst 1936) included at least three genes affecting each of the traits that differ between morphs, with thrum individuals heterozygous (*S/s*) and pin individuals homozygous (*s/s*) at the *S*-locus. However, contrary to the classic model, genomic studies in *Primula* have shown that the dominant haplotype is defined by a 278-kb region exclusively inherited by thrum individuals in hemizygosity (Li et al. 2016; Burrows and McCubbin 2017; Cocker et al. 2018). The hemizygous region harbors *CYP734A50*, a gene that simultaneously controls the position of the style (Huu et al. 2016) and female self-incompatibility (Huu et al. 2022), and *GLO*^*T*^, which determines anther position (Huu et al. 2020). The presence of paralogs of *S*-linked genes at different genomic locations suggests stepwise assembly of the genes at the *S-*locus (Huu et al. 2020; Potente et al. 2022). Subsequently, hemizygosity and *S-*linked candidate genes for style elongation have also been documented in distylous *Turnera* (Shore et al. 2019; Matzke et al. 2021).

Hemizygosity at the *Primula* and *Turnera S*-loci has important implications for our understanding of the origin, maintenance and breakdown of distyly. Although inversions are the most common types of genomic rearrangements associated with supergenes (Gutiérrez-Valencia et al. 2021), hemizygosity has been suggested to be a common feature of distyly supergenes (Kappel et al. 2017; Barrett 2019; Shore et al. 2019). Genomic and functional studies in additional distylous systems can shed light on whether distyly *S*-loci exhibit convergent genetic architecture, gene function, and evolution.

*Linum* presents a remarkable opportunity to study the evolution and loss of distyly supergenes, because it exhibits dynamic evolution of mating systems, including multiple independent origins and losses of distyly (Armbruster et al. 2006; McDill et al. 2009; Ruiz-Martín et al. 2018). Although molecular studies have led to the identification of candidate *S*-locus genes (Ushijima et al. 2012; Ushijima et al. 2015), the distyly *S*-locus in *Linum* has not been fully sequenced nor characterized even though *Linum* is one of the genera where distyly was first studied (Darwin 1863; Darwin 1877).

Here, we uncover the genetic architecture and evolution of the supergene that governs distyly in *L. tenue* (Fig. 1). By first building a chromosome-level genome assembly, we identify the *Linum S*-locus and show that it is characterized by the presence of a thrum-specific hemizygous ∼260-kb region that harbors nine predicted genes, including a candidate gene for style length. By extending the study of distyly supergenes to a novel system, we demonstrate remarkable molecular convergence in supergene architecture and evolution across widely diverged systems, and shed light on the origin and maintenance of an iconic floral polymorphism.

## Results

### A Chromosome-Level Genome Assembly of *L. tenue*

We produced a high-quality *de novo* genome assembly of a *L. tenue* thrum individual to aid the identification of the distyly *S-*locus. We first generated a partially phased contig assembly based on high-coverage (∼170x) PacBio long-read data, scaffolded with chromatin conformation capture (Hi-C) data and polished with 10x linked-reads (see Fig. S1 and Supplementary Methods, Supplementary Note). We generated a genetic map to correct and anchor primary scaffolds to 10 linkage groups (LG), corresponding to the 10 haploid chromosomes of *L. tenue* (Pastor et al. 1990). The resulting 702.1 Mb haploid genome assembly was highly contiguous (N50=69.3 Mb) and complete (Complete BUSCOs=94.2%, flow cytometry-based estimate of genome size ∼690 Mb) (Table S1, Supplementary Tables). Annotation of the assembly using both *ab initio* prediction tools and RNA-sequencing evidence led to the identification of 52,826 coding sequences (Complete BUSCOs=93.8%, based on the longest isoforms per gene; Table S2, Supplementary Note) and 595,563 non-overlapping repetitive sequences that make up ca. 49.36% of the genome.

### The Distyly Supergene is Hemizygous in Thrum Individuals

Presence-absence variation is an important feature of distyly supergenes in *Primula* and *Turnera* (Li et al. 2016; Cocker et al. 2018; Shore et al. 2019). To identify genomic regions harboring presence-absence variation between floral morphs, we analyzed genome coverage for 21 pin and 22 thrum individuals sequenced with short reads. We identified a ∼300 kb region between 38.40 and 38.70 Mb on LG10 with significantly elevated coverage in thrums relative to pins (Fig. 2a) (Wilcoxon signed-rank test followed by Bonferroni correction, *P*<0.001 for windows between LG10: 38.40-38.70 Mb, and *NS* for all remaining 50 kb windows across the entire genome; mean coverage across samples: mean=29.45, SD=9.73, *n*=43) (Fig. S2 in Supplementary Results, Supplementary Note). In contrast, there were no windows where pin individuals had significantly higher coverage than thrums. Compared to flanking regions, coverage in the 38.40-38.70 Mb region of LG10 was significantly reduced in thrum individuals, as expected if thrums are hemizygous (Fig. 2a, Fig. S2 in Supplementary Results, Supplementary Note). The presence of a ∼260 kb thrum-specific region on LG10 was confirmed by the alignment of 10x Genomics linked-reads Supernova assemblies of thrum (*n*=4) and pin (*n*=5) individuals to our *L. tenue* reference genome (Fig. S3 in Supplementary Results, Supplementary Note). The annotation of this ∼260 kb insertion-deletion (indel) showed that it harbored a gene encoding a homolog of the *L. grandiflorum* style-expressed thrum-specific protein TSS1 (see section *Distyly Candidate Genes at the S-Locus Show Thrum-Specific Expression*) suggested to be *S-*linked in *L. grandiflorum* (Ushijima et al. 2012; 2015). We therefore considered the ∼260 kb thrum-specific region on LG10 a candidate region for the distyly *S-*locus in *L. tenue*.

**Fig. 2.**
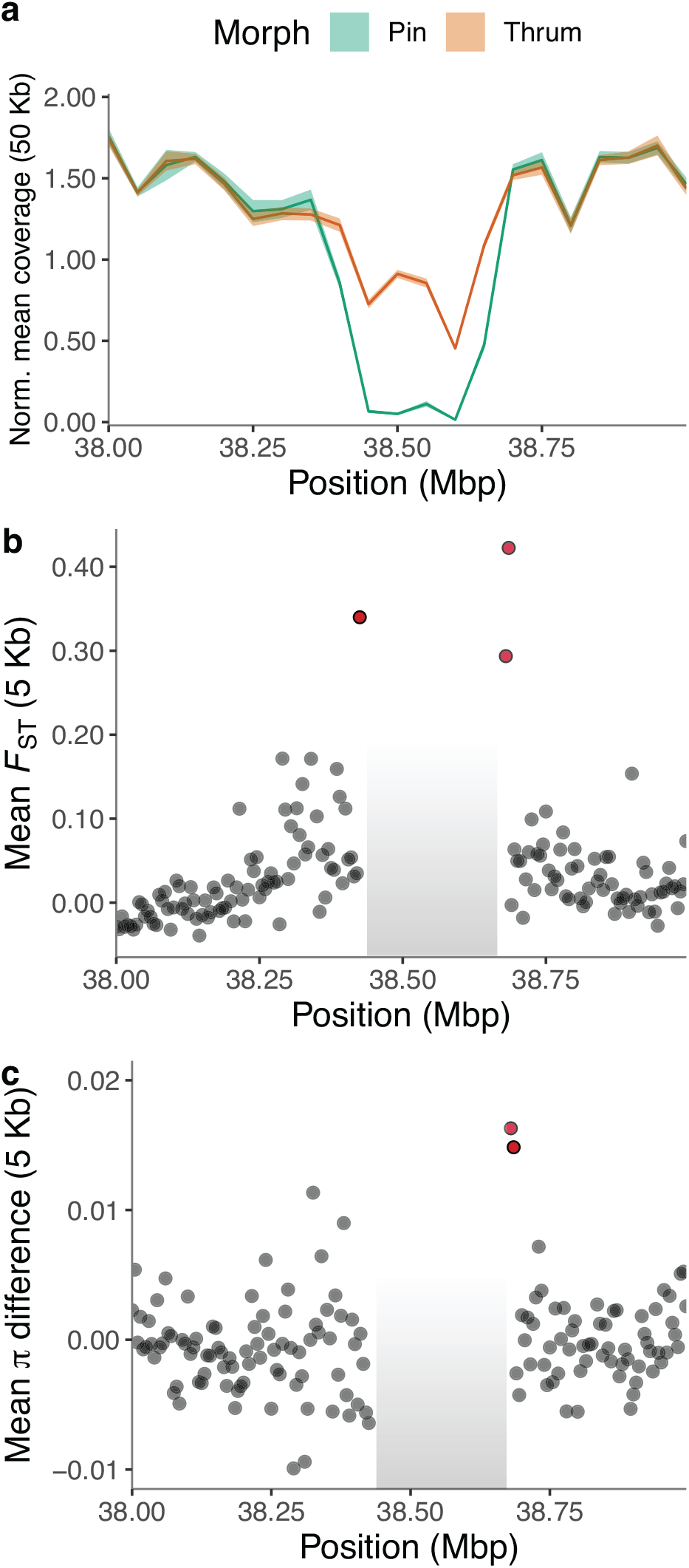
**a**, Differences in coverage between thrum (*n*=26) and pin (*n*=25) samples indicates the existence of a ∼260 kb hemizygous region in LG10 (shaded regions indicate 95% CI), as supported by the existence of six 50 kb adjacent windows that are absent in pin samples. Estimates of **b**, genetic differentiation (*F*_*ST*_) and **c**, differences in mean *π* between thrum and pin sequences. **b**-**c**, Calculations were conducted on 5 kb windows across the genome (excluding the ∼260 kb hemizygous region) using samples from population SMT (thrum=13 and pin=13) (see estimates for population CL: Fig. S4 in Supplementary Results, Supplementary Note). Windows with statistically significant estimates of F_*ST*_ and *π* distance between morphs are highlighted in red (*P* < 0.01, after 1000 replicates of permutation test per window, followed by FDR correction with the Benjamini-Hochberg method). The masked ∼260 kb hemizygous region is shaded in grey in panels b and c.

To conclusively identify the distyly supergene in *L. tenue*, we performed genome-wide association mapping (GWAS) of 8.7 million SNPs in 21 thrum and 21 pin individuals and identified a single genomic region on LG10 (positions 38,425,470-38,686,519) as significantly associated with floral morph (Fig. 3a) and thus likely to harbor the *L. tenue* distyly supergene. There were 79 SNPs immediately flanking (within 3.5 kb) the previously identified ∼260 kb hemizygous region that were strongly associated with floral morph (Fisher’s exact test, followed by FDR with the Benjamini-Hochberg procedure, *P*<0.01) (Fig. 3b, c). High genetic differentiation (*F*_ST_) between thrum and pin individuals extended approximately 15 kb outside of the hemizygous region (Fig. 2b, c and Fig. S4 in Supplementary Results, Supplementary Note) (*P*<0.01 after FDR/BH correction, permutation test, 1000 replicates per window), and 5 kb windows immediately flanking the hemizygous region had higher nucleotide diversity (*π*) in thrum than in pin individuals, as expected if these regions were heterozygous in thrum individuals (Fig. 3c) (*P*<0.01 after FDR/BH correction, permutation test, 1000 replicates per window).

**Fig. 3.**
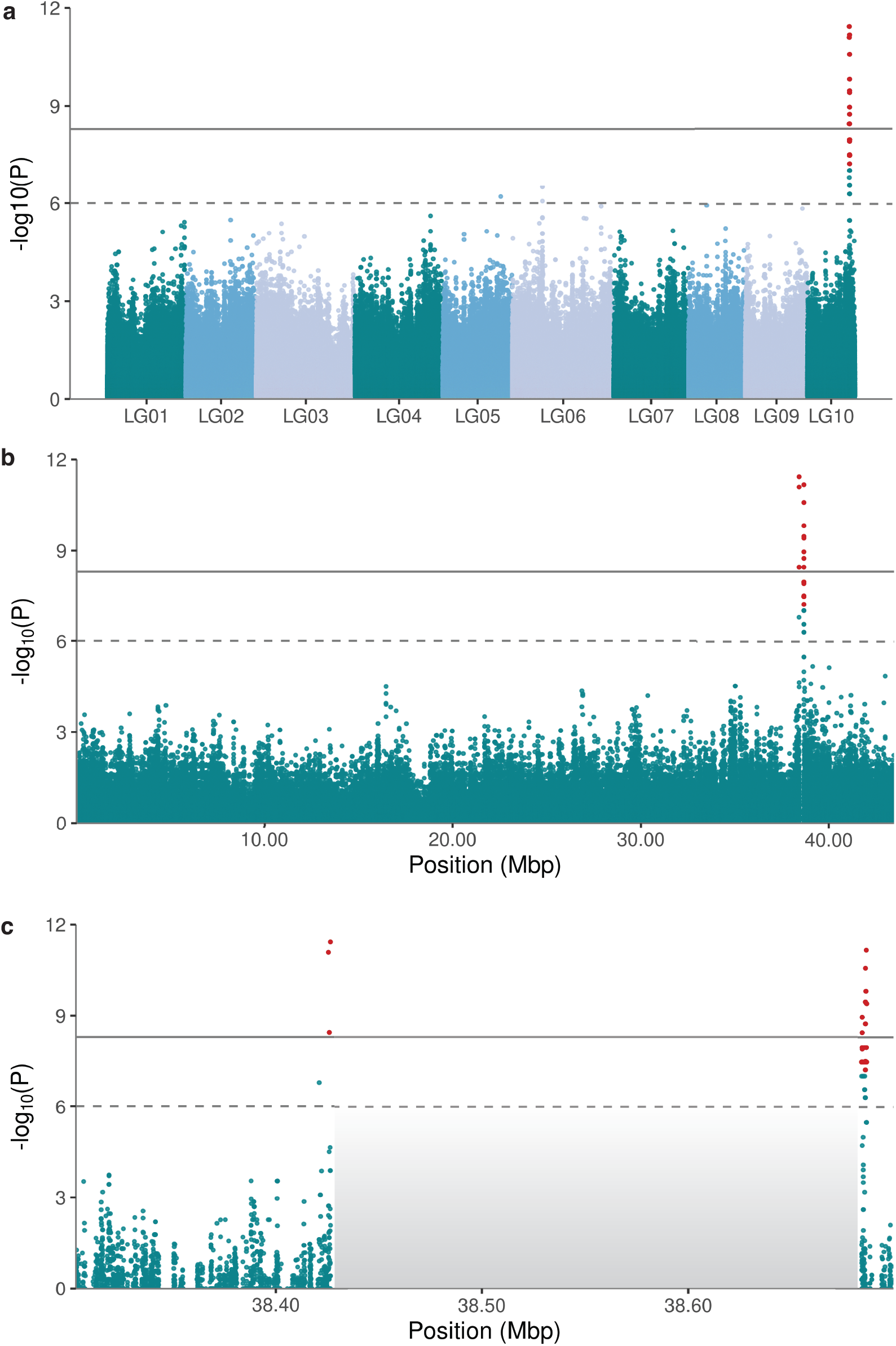
Association between SNPs genotypes and floral morph in *L. tenue* (n=42 individuals, 21 of each morph) using a GWAS analysis. Manhattan plots depict SNPs throughout **a**, the entire genome, **b**, LG10, and **c**, and loci neighboring the masked ∼260 kb hemizygous region (shaded in grey). Dashed and contiguous horizontal lines denote suggestive (-log_10_(1×10^−6^)) and significant association (-log_10_(1×10^−9^)) prior to *P* values correction. SNPs colored in red highlight loci that held significantly associated with floral morph following FDR correction using the Benjamini-Hochberg procedure, *P*<0.01.

Together, coverage analyses, GWAS and haplotype reconstruction indicate that the *L. tenue S*-locus is characterized by the presence of a ∼260 kb hemizygous region exclusive to the dominant *S* haplotype and missing from the recessive *s* haplotype. Sequence differentiation between the dominant and recessive haplotype is limited to approximately 15 kb immediately flanking the hemizygous region. Thus, the distyly *S-*locus in *L. tenue* is predominantly hemizygous in thrums (98.64% of its sequence).

### Lack of Broad-Scale Recombination Suppression Around the Distyly Supergene

The highly localized sequence differentiation between pins and thrums surrounding the hemizygous region suggests that the distyly *S-*locus is not located in a genomic region with broad-scale recombination suppression. In line with this hypothesis, map-based broad-scale recombination rates were intermediate in the *S-*locus region (∼4 cM/Mb; Fig. S5 in Supplementary Results, Supplementary Note) and linkage disequilibrium decay in natural populations (*n*=43) showed no evidence for broad recombination rate suppression outside of the hemizygous region (Fig. S6, Supplementary Note). Thus, structural variation at the *S-*locus, and not recombination suppression in a broader genomic region, is likely instrumental for maintenance of the distyly supergene in *L. tenue*.

### Evolutionary Genetic Signatures of Relaxed Selection at the Distyly Supergene

Hemizygosity of the *S-*locus in thrums can elegantly explain the absence of recombination at distyly supergenes. Morph-limited inheritance and lack of recombination could lead to genetic degeneration and accumulation of repetitive elements, but decay can be slowed down by hemizygous exposure of recessive deleterious alleles to selection (Gutiérrez-Valencia et al. 2021).

First, we compared repetitive element content at the *S*-locus compared to its flanking regions and the rest of LG10. We found that the distyly *S*-locus was enriched in TEs relative to its immediate genomic neighborhood (Fig. 4c) (*S*-locus proportion TEs in 25 kb windows: median=62.50%, *n*=9; flanking regions: median=31.8%, *n*=18) (Wilcoxon rank-sum test, *P* < 0.001) and compared to other windows in LG10 (Fig. 4b). Accumulation of TE insertions in distyly supergenes might be facilitated by the joint action of lack of recombination and reduced effective population size.

**Fig. 4.**
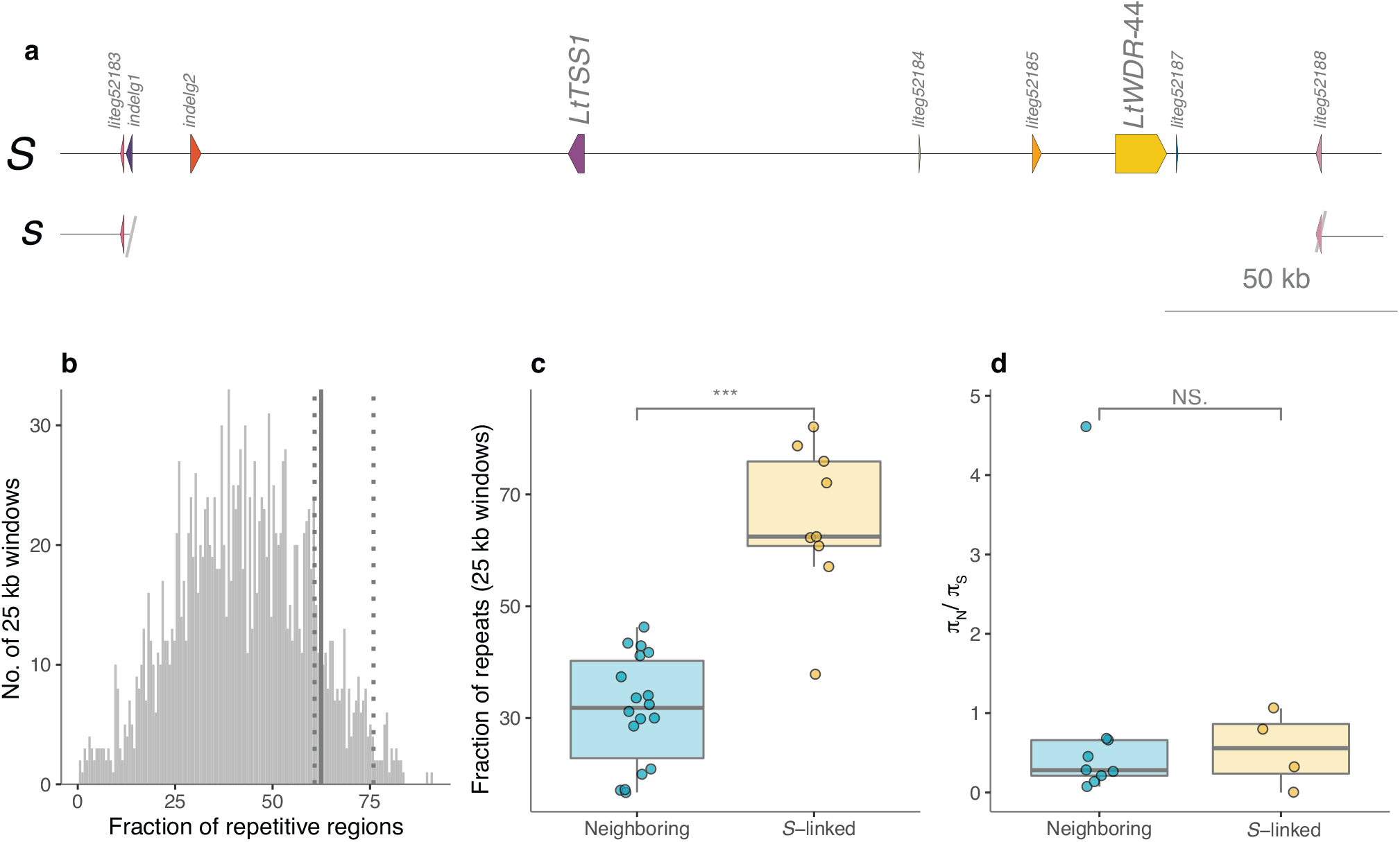
**a**, Schematic representation of the dominant *S* and recessive *s* alleles of the distyly *S*-locus, indicating the location of the two distyly candidate genes *LtTSS1* and *LtWDR-44*, only present in the dominant *S* allele. **b**, Distribution of the fraction of repetitive regions in 25 kb windows across LG10. The median value of the fraction of repeats in 25 kb windows (*n=*9) for the S-locus falls in the third quartile of the distribution (solid line represents the median=62.50%; dotted lines represent the first and third quartiles=60.77 and 75.90%). **c**, Comparison of the fraction of repetitive sequences linked to the S-locus (*n=*9 25 kb windows) and neighboring loci (*n=*9 25 kb windows for each up- and downstream regions). **d**, Comparison of π_*N*_/π_*S*_ estimates between S-locus-linked (n=4 genes with π_*s*_> 0) and neighbouring genes (n=8 genes with π_*s*_ > 0 on both sides of the *S*-locus) (*post hoc* average π_*N*_/π_*s*_: *S*-locus=0.597, neighbouring=0.300, 13 individuals from population SMT) (\**P* < 0.05, \*\**P* < 0.01, \*\*\**P* < 0.001, Wilcoxon rank sum test).

Next, using polymorphism data from thrum samples, we tested for a signature of relaxed purifying selection and found no significant difference in the ratio of nonsynonymous to synonymous polymorphism (π_*N*_/π_*S*_) between *S*-linked and neighboring genes (Fig. 4e). These results suggest that *S*-linked genes are not accumulating deleterious mutations, consistent with recent simulations showing that hemizygosity at supergenes might slow this process (Gutiérrez-Valencia et al. 2021).

### Distyly Candidate Genes at the S-Locus Show Thrum-Specific Expression

As a basis for further functional and evolutionary genetic studies, we conducted detailed annotation of the *S-*locus and analyzed RNA-eq data from floral buds, individual mature floral organs (pistils, stamens and petals), and leaves to delineate candidate genes with floral organ-specific or biased expression between morphs.

There were nine protein-coding genes in the dominant *S-*haplotype, excluding six genes with TE-related functions (Fig. 4a; Table S3, Supplementary Tables). Five of the *S-*locus genes had no known function (Table S3, Supplementary Tables). Genes encoding proteins with assigned functions included: a protein of the VASCULAR-RELATED UNKNOWN PROTEIN1 (VUP1) family involved in regulating tissue growth modulated by hormone signaling (see discussion of *LtTSS1* below), an amino acid binding protein (WD repeat-containing protein 44, WDR44), a tetratricopeptide repeat-containing protein and a gene encoding a protein belonging to a gene family that includes genes with functions in pollen development (PANTHER family: PTHR34190, GO-term: GO:0009555) (Table S3, Supplementary Tables). The latter gene (LITEG00000052183) was the only *S-*locus gene that was also present in the recessive *s-*haplotype, located approximately 1 kb downstream of the ∼260 kb indel (Fig. 4a). At the upstream limit of the indel, we further annotated a complete gene (LITEG00000052188) in the dominant *S* haplotype that was truncated in the recessive *s* haplotype (Fig. 4a).

The dominant *S-*haplotype harbored a strong candidate gene for style length, *L. tenue TSS1* (*LtTSS1*). This gene shows homology to the *L. grandiflorum* gene *TSS1* (significant alignment with Blastp: E-value=3×10^−6^, sequence identity=43%, length=173, gaps=8%) (Fig. 4a; Table S4, Supplementary Tables), a thrum-specific *S-*linked gene in *L. grandiflorum* (Ushijima et al. 2015). *LtTSS1* had strongly elevated and pistil-specific expression in thrum relative to pin pistils and buds (log2FC>10; Fig. 5a, b). The amino acid sequence of *LtTSS1* shows high similarity to a protein of the VUP1 family (best match=IPR039280, E-value=7.8×10^−11^, percent identity=43.4%) (Blast search with UniProt, The UniProt Consortium 2021). In *A. thaliana*, overexpression of *VUP1* suppresses cell expansion leading to reductions in organ size, including shorter floral organs (Grienenberger and Douglas 2014). As cell length is shorter in thrum than pin styles of both *L. tenue* and *L. grandiflorum* (Ushijima et al. 2015; Foroozani 2018, in prep), *TSS1* is a strong candidate for a locus regulating style length.

**Fig. 5.**
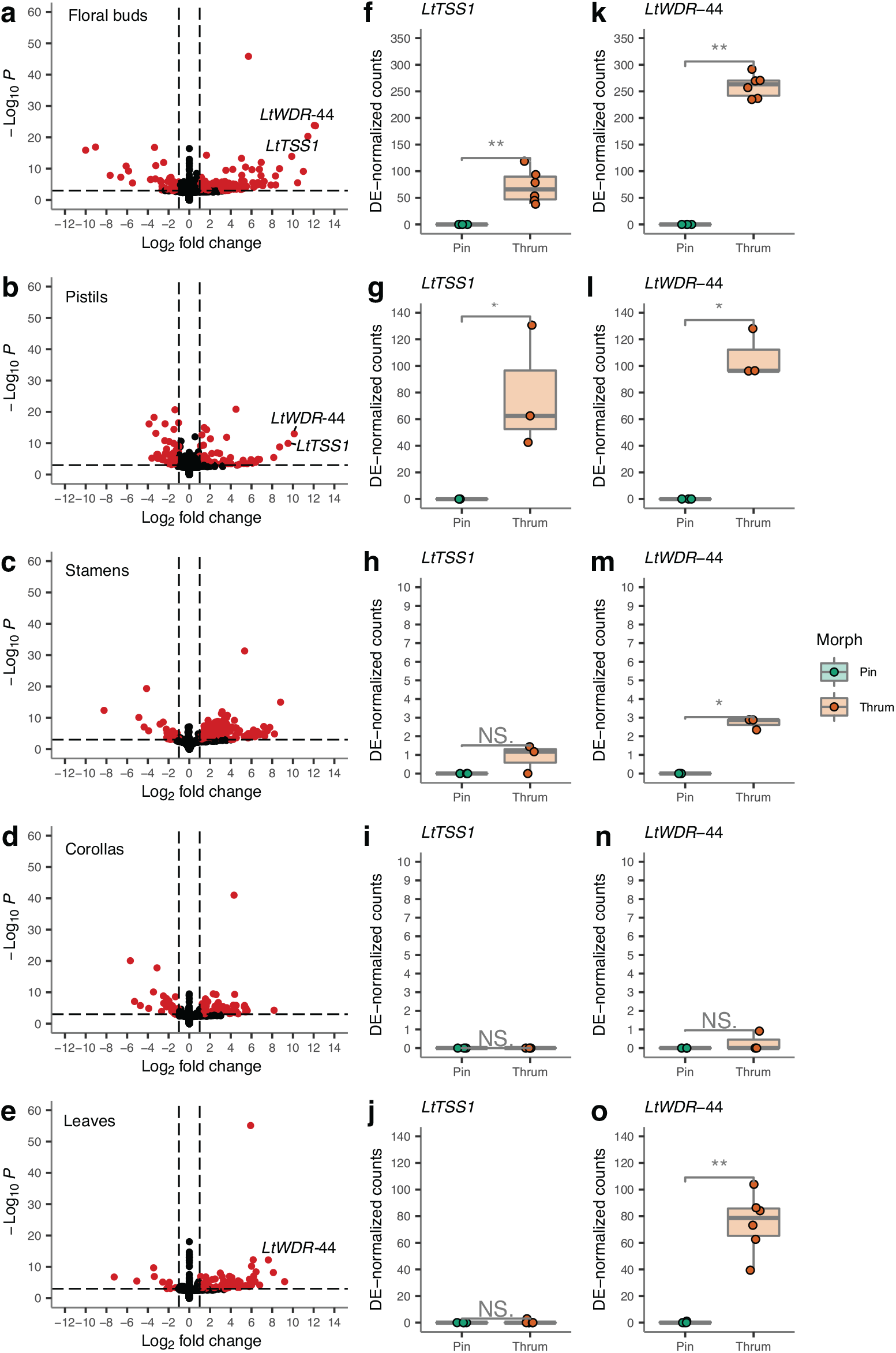
Differential expression in thrum and pin floral organs and leaves, including **a**, entire floral buds (n thrum=6, pin=4), **b**, pistils (n thrum=3, pin=3), **c**, stamens (n thrum=3, n pin=3), **d**, corollas (n thrum=3, n pin=3) and **e**, leaves (n thrum=6, pin=4). Differentially expressed genes (DEG) are highlighted in red (horizontal dashed line: cut-off for the adjusted *P*-value < 0.001, vertical dashed lines: Log2 fold change > |1.5|). Since *LtTSS1* and *LtWDR-44* were identified as DEG upregulated in thrum floral buds and pistils, we compared normalized counts of both genes between pin and thrum samples in **f, k**, floral buds, **g, l**, pistils, **h, m**, stamens, **i, n**, corollas and **j, o**, leaves (\**P* < 0.05, \*\**P* < 0.01, Wilcoxon rank sum test).

The *S-*linked gene *LtWDR-44* is an interesting candidate gene for floral differentiation between morphs due to its annotation suggesting functions in developmental growth and hormone- mediated signaling. *LtWDR-44* also had higher expression in thrums than pins in buds, pistils, stamens and leaves (Fig. 5a, b, e). *LtWDR-44* was primarily but not exclusively expressed in pistils (median: pistils=3.72 TPM, stamen=0.13 TPM) (Tables S4 and S5, Supplementary Tables). Three additional *S*-linked genes (INDELG00000000001, LITEG00000052183 and LITEG00000052188) were expressed both in petals and stamens, and are potential candidate genes for anther position or pollen specificity, respectively, whereas the remaining *S*-linked genes were not detectably expressed in our samples. These results help delineate candidate genes at the *S-*locus, confirm thrum-specific expression *S-*linked genes, and demonstrate that the style length candidate gene *LtTSS1* has pistil-specific expression in *L. tenue*.

### Cell Wall-Related Genes are Differentially Regulated in Thrum and Pin Pistils and Stamens

To investigate sets of genes and pathways that might be regulated by *S-*linked genes, we performed differential expression and gene set enrichment analyses. We were specifically interested in differential expression of cell wall-related genes between floral morphs, as altered expression of such genes is an effect of overexpression of the *TSS1* homolog *VUP1* (Grienenberger and Douglas 2014).

Floral buds showed a higher proportion of differentially expressed genes (DEG) between morphs compared to leaves, and among mature floral organs, stamens had the largest fraction of differentially expressed genes, followed by pistils and petals (Table 1). Cell wall-related genes (GO:0071555; GO:0042545) were significantly enriched among DEG (*P*≤ 0.05; Table S6, Supplementary Tables) in both pistils and stamens, which differ in length and cell size between floral morphs (Foroozani 2018, in prep), as well as in buds, but not in petals and leaves (Table S6, Supplementary Tables). In pistils, GO terms showed significant enrichment in cell wall organization through oxido-reductive activity mainly located in the cell walls and anchored to membrane compartments. Thrum-specific expression of *LtTSS1* in pistils is thus associated with altered regulation of genes involved in cell wall organization, potentially resulting in shorter styles in thrum flowers.

**Table 1.** Summary of differential expression analysis. Significant differentially expressed genes (adjusted *P*<0.01) were classified as up- (LFC > 0) or down-regulated (LFC < 0). The analyzed sets exclude genes with zero counts across all samples, extreme count outliers (detected by Cook’s distance) and low mean normalized counts (see Supplementary Note for details).

| Tissue | Analyzed | Significant DEG (%) | LFC > 0 (%) | LFC < 0 (%) |
| --- | --- | --- | --- | --- |
| Bud | 37,001 | 123 (0.33) | 73 (0.20) | 50 (0.14) |
| Pistil | 28,615 | 101 (0.35) | 32 (0.11) | 69 (0.24) |
| Stamen | 26,908 | 223 (0.83) | 200 (0.74) | 23 (0.09) |
| Petal | 23,803 | 52 (0.22) | 23 (0.10) | 29 (0.12) |
| Leaf | 30,439 | 54 (0.18) | 37 (0.12) | 17 (0.06) |

### Evidence for stepwise assembly of the *Linum S*-locus gene set

We investigated the genomic position of paralogs of *S*-linked genes using our chromosome-level genome assembly. If the distyly supergene formed via segmental duplication, most paralogs of *S*-linked genes should stem from the same genomic region, but if they are scattered across the genome, this would support stepwise assembly of the *S*-linked gene set. Reciprocal Blast analyses identified putative paralogs of three *S-*linked genes (*LITEG00000052184, LITEG00000052185* and *LtWDR-44*) in two regions of LG1 separated by > 60 Mb, with > 2 Mb between genes at the upstream end of the chromosome (Table S8, Supplementary Tables). The closest putative paralogs of the *S-*locus genes *LITEG00000052183* and *LITEG00000052188* were found on LG9 separated by 33 Mb. We calculated synonymous divergence (*d*_*S*_) between *S*-linked genes and their putative paralogs as a proxy for the time since gene duplication (Table S8, Supplementary Tables). Our estimates varied greatly, suggesting that these genes duplicated at different times, with *LtWDR*-44 likely duplicating first (Fig. S13a, Supplementary Note).

Finally, to investigate if the scattered distribution is ancestral, we conducted similar analyses based on the high-quality reference genomes of two outgroups within the Malphigiales, cassava (*Manihot esculenta*) (Bredeson et al. 2016) and poplar (*Populus trichocarpa*) (Tuskan et al. 2006). There were four *S*-linked genes with significant matches in both genomes (Table S9, Supplementary Tables). While most genes were retrieved as located on different chromosomes, matches to *LtTSS1* and *LtWDR-44* were co-located in both genomes, although more than 0.8 and 1.5 Mb apart in the case of cassava and poplar, respectively. Taken together, these results suggest stepwise assembly of the gene set at the *Linum S*-locus (Fig. S13b, Supplementary Note).

## Discussion

Complex phenotypic polymorphisms have long fascinated biologists, yet until very recently the genetic architecture and evolution of supergenes remained uncharacterized. Here, we expand detailed characterization of distyly supergenes to *Linum*, a classic system for the study of distyly (Darwin 1863). Building on a chromosome-level genome assembly of *L. tenue*, we identify the distyly supergene and show that the dominant *S-*haplotype harbors a ∼260 kb thrum-specific region. We show that, although the dominant allele has accumulated TEs, *S-*linked genes do not show signatures of relaxed purifying selection. Finally, we characterized differential expression of *S-*linked genes potentially controlling floral organ elongation. Our results show that independently evolved *Linum, Primula*, and *Turnera S*-loci, which are the distyly supergenes studied in most detail so far, exhibit remarkable convergence in their genetic architecture with hemizygosity in thrum individuals as a shared feature.

Hemizygosity in thrums could elegantly explain both the lack of recombination and dominance of the thrum *S-*locus allele, but structural variation could potentially constitute either a cause or a consequence of suppressed recombination. In *Primula*, the location of the *S*-locus in a pericentromeric region (Li et al. 2015; Kappel et al. 2017; Potente et al. 2022) might have favored supergene formation if recombination rates were ancestrally low in this region. In contrast, at the *Linum* distyly supergene, we observed localized recombination suppression largely coinciding with the boundaries of the indel, without broad-scale recombination suppression surrounding the supergene. This observation strongly suggests that structural variation is a likely cause and not a consequence of suppressed recombination.

The joint effects of suppressed recombination and morph-limited inheritance can precipitate accumulation of deleterious mutations. This process is expected to be slower for supergenes harboring indels than inversions, because recessive deleterious mutations in hemizygous regions are directly exposed to selection (Gutiérrez-Valencia et al. 2021). At the *Linum* distyly supergene, we observed an enrichment of TEs, but no signature of relaxed purifying selection on *S-*linked genes. This result mirrors recent findings in *Primula* (Cocker et al. 2018; Potente et al. 2022) and while efficient selection in hemizygous regions could contribute to this pattern, sequence constraints on *S-*linked genes resulting from their role in distyly could also be involved.

How then did the *Linum* distyly supergene originate? While detailed studies in a comparative framework will be required to conclusively answer this question, our results provide some information on the likely age and sequence of events. The finding that the *L. tenue S-*locus harbors *LtTSS1*, an ortholog of the *L. grandiflorum* thrum-specific and *S*-linked gene *TSS1* (Ushijima et al. 2012; Ushijima et al. 2015) suggests that presence-absence polymorphism at the *S-*locus could have been retained for at least 30-40 million years, since the divergence of *L. tenue* and *L. grandiflorum* (Ruiz-Martín et al. 2018; Maguilla et al. 2021). Furthermore, our finding that paralogs of *S-*linked genes are found in disparate genomic locations both in *L. tenue* and in outgroups suggests that the *S-*locus gene set evolved in a stepwise process involving several gene duplications and translocations, similar to the *Primula S-*locus (Huu et al. 2020; Potente et al. 2022). These results imply that the presence-absence polymorphism at the *Linum* distyly *S-*locus likely did not originate through one large segmental duplication.

The independent evolution of distyly supergenes with hemizygosity in thrums inevitably raises the question: why did evolution repeatedly favor this genetic architecture? One possibility is that hemizygosity facilitates establishment of new advantageous mutations, as they will be directly exposed to efficient selection (Haldane 1927). Under the “selfing avoidance” model, the first mutation would have led to the origin of pollen incompatibility in a monomorphic and self-compatible population (Charlesworth and Charlesworth 1979). In contrast, the “pollen transfer” hypothesis suggests that the first mutation would modify the length of the style in a population of approach herkogamous flowers (Lloyd and Webb 1992). Although it is unknown how distyly evolved in *Linum*, ancestral state reconstruction studies suggest several transitions from approach herkogamy to distyly (Ruiz-Martín et al. 2018). Determining whether mutations affecting pistil length were the first step in the formation of the *Linum* distyly *S*-locus will shed further light on this question.

A particularly strong candidate gene for the control of style elongation in *L. tenue* is the *S-* linked gene *LtTSS1*, due to its pistil-specific expression only in thrums and inferred biological function. Previous morphometric work in *Linum* reported longer style cells in pin compared to thrum flowers, indicating that style length dimorphism is caused by differences in cell elongation (Ushijima et al. 2015; Foroozani 2018, in prep). Importantly, because *LtTSS1* significantly resembles genes from the VUP1 family, which have been linked to organ size reduction through repressed cell expansion (Grienenberger and Douglas 2014), it is likely that the expression of *LtTSS1* in thrum pistils limits cell expansion leading to shorter styles. The expression of *LtTSS1* in floral buds suggests the early onset of developmental processes underlying differences in style length between morphs, and the significant enrichment of genes involved in cell wall modification among floral bud and pistil DEG between morphs further supports this idea. Cell length reduction leading to shorter thrum pistils is controlled by hemizygous *CYP734A50* in *Primula* (Huu et al. 2016) and *TsBAHD* in *Turnera* (Shore et al. 2019; Matzke et al. 2021). Interestingly, both genes encode brassinosteroid-inactivating enzymes (Huu et al. 2016; Matzke et al. 2021), which not only determine pistil length but also female incompatibility function (Matzke et al. 2021; Huu et al. 2022). Heteromorphic self-incompatibility has been documented in *L. tenue* (Murray 1986), and as overexpression of *VUP1* down-regulates the expression of brassinosteroid-responsive genes (Grienenberger and Douglas 2014), it will be relevant to investigate whether pistil length and female incompatibility might also have evolved through alterations in the brassinosteroid pathway in *Linum*. Further functional studies of *LtTSS1* and other *S*-linked genes that we have identified here will be required to conclusively elucidate the genetic and developmental mechanisms underlying distyly in *Linum*.

Taken together, our findings in combination with those of earlier studies indicate remarkable convergence with respect to the genetic architecture, origin and evolution of distyly supergenes in *Linum, Primula* (Li et al. 2016; Cocker et al. 2018; Huu et al. 2020; Potente et al. 2022) and *Turnera* (Shore et al. 2019), and suggest that convergence may extend to the level of molecular pathways controlling distyly. Our chromosome-level genome assembly and detailed characterization of the *Linum* distyly *S-*locus augment our knowledge on the evolution of *S*-loci across distylous lineages and motivate further work to better understand the conditions favoring similar mechanisms and processes for the origin and maintenance of supergenes.

## Methods

### Genome Assembly

We produced a high-quality *de novo* genome assembly of a single outbred *L. tenue* thrum individual based on high-coverage (∼170x) PacBio long-read data (V3.0 chemistry on Sequel), with scaffolding using Hi-C data and a genetic linkage map. We first generated a partially phased diploid assembly using FALCON and FALCON Unzip (Chin et al. 2016) that was polished using Arrow. The resulting primary assembly was 879.49 Mb (No. sequences=1222, contig N50=1.43 Mb; Table S1, Supplementary Tables) and had an associated set of haplotigs of 161.13 Mb (No. sequences=1540). We used PurgeHaplotigs (Roach et al. 2018) to reassign misclassified primary contigs as haplotigs based on read-depth analyses and repeat annotations (Fig. S1 in Supplementary Methods, Supplementary Note).

We next scaffolded the assembly using chromatin conformation and capture (Hi-C) data (Lieberman-Aiden et al. 2009) generated using the Proximo Hi-C 2.0 protocol of Phase Genomics (Seattle, WA). The purged version of the primary assembly (615.74 Mb, No. sequences=553), the set of haplotigs originally identified with FALCON-Unzip concatenated to those reassigned by PurgeHaplotigs (420.59 Mb, No. sequences=2,128), and the Hi-C data were used to phase and scaffold the genome assembly. First, the partially phased PacBio-based assembly was processed together with the Hi-C data using FALCON-Phase v1.2.0 (Kronenberg et al. 2021) available as the pb-assembly package (v0.0.8) from Bioconda (https://bioconda.github.io/recipes/pb-assembly/) to produce two full length pseudo-haplotypes for phase0 and 1. For scaffolding, Hi-C reads were aligned to phase0 of the FALCON-Phase assembly using BWA-MEM (v0.7.17) (Li and Durbin 2009) followed by filtering with SAMtools (v1.9) (Li et al. 2009) and the script PreprocessSAMs.pl (https://github.com/tangerzhang/ALLHiC/blob/master/scripts/) to keep only links with strong signals in the Hi-C data set, and the scaffolds of phase0 were chained into ten pseudochromosomes using ALLHiC (v0.9.13) (Zhang et al. 2019). This step was followed by a rescue step to assign unplaced contigs into partitioned clusters as implemented in ALLHiC (v0.9.13) (Zhang et al. 2019). After a second round of FALCON-Phase using the scaffolded version of phase0 as primary assembly and the non-scaffolded version of phase1 as associated contigs, misplaced scaffolds were reassigned to each phase. This yielded a chromosome-scale (scaffold N50 64.2 Mb) phased pseudo-diploid assembly of ten pseudochromosomes totaling 703.83 Mb for phase0 and 702.16 Mb for phase1 (Table S1, Supplementary Tables). To polish the assembly, 10x Genomics linked reads were aligned to each phase of the reference using Long Ranger (https://github.com/10xGenomics/longranger), followed by two rounds of Pilon (Walker et al. 2014). We further edited phase1 to retain a region (contig 000228F_arrow) preliminarily identified as associated with floral morph (Supplementary Methods, Supplementary Note), that was removed during the first step of the Hi-C scaffolding procedure (Supplementary Methods, Fig. S7 and S8, Supplementary Note). Assembly statistics for each step of the assembly were obtained using the script gaas_fasta_statistics.pl (https://github.com/NBISweden/GAAS/tree/master/bin).

We anchored scaffolds to pseudochromosomes and corrected mis-joins using Lep-Anchor (Rastas 2020). As Lep-Anchor requires a haploid assembly, we processed a haploid assembly representing one of our phases of our scaffolded pseudo-diploid assembly (phase1). First, we generated a linkage map in Lep-MAP3 (Rastas 2017) based on short-read sequencing data from a mapping population of 180 F2 individuals (see Supplementary Methods in Supplementary Note). Second, we generated a chains file made in HaploMerger (Huang et al. 2012). Third, we generated a paf file mapping the FALCON phase round 1 assembly to the scaffolded pseudo-diploid assembly (phase1) with minimap2 (Li 2018). We input these data in Lep-Anchor to obtain 10 pseudochromosomes.

Inspection of Lep-Anchor Marey maps identified two regions, LG1:10,941,743-18,235,911 and LG5:1-2,929,725 (Supplementary Methods and Fig. S9, Supplementary Note), where the recombination map position and the physical position of the marker did not match. Further inspection of the Hi-C data based on a HiGlass visualization (Kerpedjiev et al. 2018) supported this conclusion, and thus we manually edited these two regions. In the first case we moved the region to the end of LG1 and in the second we inserted it at position 56,376,345 of LG5. The final haploid genome assembly contained 10 pseudochromosomes with a total length of 702.08 Mb and was highly contiguous (scaffold N50 69.33 Mb) and complete (BUSCO complete 94.2%) (Table S1, Supplementary Tables).

### S-locus Haplotype Phasing

To phase haplotypes at the *S-*locus we mapped 10x Genomics linked-read sequencing data from thrum individual SMT-34-3 to the reference genome using Long Ranger (https://github.com/10xGenomics/longranger) which through FreeBayes (Garrison and Marth 2012) calls phased variants. Within the region identified as associated with floral morph in GWAS analyses, we verified the presence of two divergent haplotypes differing with regard to an indel, a ∼260 kb region (LG10:38,426,515-38,684,012) within a 1 Mb phased block (LG10:37,829,718-38,865,685). The haplotype information was used to modify the reference genome sequence in this region to represent the dominant *S-*haplotype prior to further analyses (Figs. S11, S12 and Supplementary Methods, Supplementary Note).

### Genome Annotation

The identification of coding sequences relied on the usage of curated and custom protein sequences databases, and the assembly of the transcriptomes from different tissues. We used the Uniprot Swiss-Prot database (Magrane and Consortium 2011) to produce a non-redundant protein sequence database that consisted of 563,972 proteins (downloaded on December of 2020). We also identified and downloaded taxon-specific protein data bases from UniProt and Ensembl, including the proteomes of Rosids (rosids_swissprot.fasta with 22,046 proteins, and Vitis_vinifera.12X.pep.all.fa, with 29,927 proteins), the order Malpighiales (Populus_trichocarpa.Pop_tri_v3.pep.all.fa with 73,012 proteins) and the genus *Linum* (Lusitatissimum_200_v1.0.protein.fa with 43,484 proteins).

We generated *de novo* transcriptome assemblies for annotation using RNA-Seq data from mature flowers, floral buds, leaves and stems from two thrum individuals. After error correction, adapter and quality trimming and removal of ribosomal reads (Supplementary Methods in Supplementary Note), we generated *de novo* assemblies with Trinity (2.9.1) using default settings (Grabherr et al. 2011; Haas et al. 2013). Additional assemblies were produced by processing the *in silico* normalized files created through Trinity with Velvet (Zerbino and Birney 2008) and Oases (Schulz et al. 2012). The transcriptomes obtained with the pipelines of Trinity and Velvet/Oases were merged using EvidentialGene (http://eugenes.org/EvidentialGene/) to identify the primary transcripts and alternate transcripts/isoforms accepted as valid. In addition, we also used a reference-guided assembly approach. RNA-Seq data and the chromosome-level reference genome were processed through the pipeline TranscriptAssembly (https://github.com/NBISweden/pipelines-nextflow/tree/master) that uses fastp (Chen et al. 2018) for quality control and preprocessing of raw files, HISAT2 for the alignment of RNA reads (Kim et al. 2015), and StringTie (Pertea et al. 2015) to assembly transcripts from each tissue independently.

To identify repeats and mask the assembly prior to structural annotation, we created a custom repeat library modelled using RepeatModeler (Smit and Hubley 2008) and RepeatMasker (Smit et al. 2013). Since protein-coding genes can contain repetitive sequences, the library of repeats was vetted against the protein set (after transposon removal) to exclude any nucleotide motif present in low-complexity coding sequences. The final identification of repetitive sequences in the genome was conducted using RepeatMasker (Smit et al. 2013) and RepeatRunner (Smith et al. 2007), allowing the identification of highly divergent repeats and protein coding portions.

Gene models were constructed using MAKER (Holt and Yandell 2011) guided by evidence from both aligned transcript sequences and reference proteins, and were then used to train *ab initio* prediction tools (https://github.com/NBISweden/pipelines-nextflow/tree/master/AbinitioTraining). Once the *ab initio* tools were trained, a new run of MAKER was conducted. Two prediction strategies were conducted and gene models were compared (see details in Supplementary Methods, Supplementary Note). The approach using only Augustus (Stanke et al. 2008) lead to a higher percentage of genes identified compared with the Augustus + Snap (Korf 2004) gene build (93.8 vs 91.6% BUSCO v4.0.2 complete, respectively) (Table S2, Supplementary Tables), and the visual inspection of the results showed a clear reduction in false positive prediction. Hence, the Augustus only-based was retained for further analyses.

We manually annotated the *S-*locus region (LG10: 38,426,515-38,684,012) which was of particular interest for this study. Specifically, we first predicted gene models using Augustus (Stanke et al. 2008) with softmasking=0 and then sought additional evidence supporting the existence of the predicted protein coding sequences from our RNA sequencing data (Supplementary Methods, Supplementary Note). If a certain protein coding sequence was present both in the automatic and manual annotations, we kept only one of them for downstream analyses.

Functional annotation of the translated CDS features was performed using the pipeline FunctionalAnnotation (https://github.com/NBISweden/pipelines-nextflow/tree/master/FunctionalAnnotation). This pipeline uses Blast and InterProScan (Hunter et al. 2012) to retrieve information on protein function from 20 different sources, which was associated to each mRNA feature. To infer gene and protein names, protein sequences were blasted against the Uniprot/Swissprot reference data set, and hits with the best score (E-values < 1×10^−6^) were kept.

The annotation of tRNA sequences relied on tRNAscan (1.3.1) (Lowe and Eddy 1997) followed by the removal of features with AED scores equal to 1. Finally, other ncRNAs were predicted using the database Rfam (Nawrocki et al. 2015) using only highly conserved eukaryotic ncRNA families, and the co-variance models provided by Rfam were then processed using Infernal (Nawrocki and Eddy 2013).

### Population Genomic Analyses

To identify regions with coverage differences between thrum and pin individuals, we analyzed whole-genome sequences (Illumina short reads) from 43 individuals with known morph from two populations (Supplementary Methods, Supplementary Note; Table S7, Supplementary Tables). After adapter and quality trimming using ‘bbduk’ from BBMap/BBTools (Bushnell 2015), reads were mapped to the chromosome-scale assembly with BWA-MEM (v0.7.17) and duplicated reads were removed using MarkDuplicates from Picard tools v2.0.1 (Broad Institute 2019). Sites in repetitive regions were identified using RepeatMasker (Smit et al. 2013) and removed from consideration. We estimated per-window genome coverage in 50 kb windows separately for pin and thrum samples using BEDTools (Quinlan and Hall 2010). We tested for a difference in median coverage between thrum and pin samples for each 50 kb window using a two-sided Wilcoxon signed-rank test followed by Bonferroni correction.

We estimated polymorphism and divergence using the same short-read data as for coverage analyses. BAM files were processed with SAMtools/bcftools (Danecek and McCarthy 2017) to produce genotype likelihoods from sequence alignments using the function ‘mpileup’, and the VCF file containing variants (SNPs/INDELs) and invariant sites was created with the function ‘call’ (see Supplementary Methods in the Supplementary Note). The VCF file was further processed to only keep biallelic SNPs and invariant sites, and filtered based on quality (QUAL > 20), coverage (AVG(FMT/DP) > 10 & AVG(FMT/DP) < 50) and data missingness (F_MISSING < 0.2) using the function filter from bcftools. We removed repetitive regions and excluded the thrum-specific hemizygous region (LG10 38,426,515-38,684,012) prior to estimating polymorphism and differentiation/divergence statistics (Supplementary Methods in Supplementary Note). The resulting repeat-masked VCF file was processed with pixy (Korunes and Samuk 2021) to estimate F_*ST*_ (between thrum and pin samples) and *π* in thrum and pin samples in 5 kb windows. Each population (CL=17 and SMT=26 samples) was analyzed independently, and sample-morph association files were provided to pixy in the argument ‘--populations’. To identify which windows show significant F_*ST*_ and differential *π* between morphs, we conducted permutation tests in R. Approximate test-statistic distributions for each window were obtained by permuting the list of sample ID – morph association 1000 times, and the observed value was compared with the distributions to calculate the *P*-values, followed by adjustment for multiple testing with the FDR method using the Benjamini-Hochberg approach.

### Recombination Rate and Linkage Disequilibrium Estimates

We converted the genetic map obtained from Lep-MAP3 (Supplementary Methods in Supplementary Note) in the Rqtl format (Broman et al. 2003). After polarization and filtering to retain markers with monotonically increasing genetic distance with physical distance, the genetic map contained 2,471 markers. The recombination rate for each chromosome was calculated with the ‘est.recrate’ function of the xoi R package (https://github.com/kbroman/xoi). Linkage disequilibrium decay estimates were obtained using biallelic SNPs previously identified using data from both populations SMT and CL (*n=*43) as described in *Population Genomic Analyses*. LD decay analyses were performed with ngsLD (Fox et al. 2019) calculating the LD on SNPs with a minor allele frequency higher than 0.1 and at a maximum distance of 20 kb between each other. These estimates of r^2^ decay were obtained for the ∼260 kb indel, three down- and three upstream 250 kb windows neighboring the region, and all LG10-linked windows to obtain a mean estimate.

### Genome-Wide Association Mapping

We conducted a GWAS to identify loci associated with floral morph. Only biallelic SNPs derived from the VCF file produced in *Population Genomic Analyses* were included. We tested for an association between morph and SNP genotype in PLINK (v1.90b4.9) (www.cog-genomics.org/plink/1.9/, Chang et al. 2015) using Fisher’s exact test, with significance adjustment using the Benjamini-Hochberg procedure to control the False Discovery Rate (FDR) (Benjamini and Hochberg 1995) (Supplementary Methods in Supplementary Note).

### Comparison of TE content between the distyly *S*-locus and neighboring windows

We estimated and compared the percentage of TEs between *S-*linked and neighboring regions. For this, we relied on the chromosome-level genome assembly and its corresponding annotation of repetitive elements (Supplementary Methods in Supplementary Note). Microsatellites (STRs and simple repeats) and regions of low complexity were not considered in these analyses, and the fraction of TEs in 25 kb windows were compared between the *S*-locus and the neighboring regions using a Wilcoxon signed-rank test. Finally, the annotation of repetitive elements was summarized to characterize the most abundant types of TEs (DNA transposons, LINE, LTR, RC or other) for all LG, LG10 and the *S*-locus (Supplementary Methods in Supplementary Note).

### Comparison of π_N_/π_S_ between distyly *S*-locus-linked and neighboring genes

To investigate if π_*N*_*/*π_*S*_ estimates differ between distyly *S*-locus-linked and neighboring genes, we produced a thrum-only VCF file containing biallelic SNPs and invariant sites (SMT=13 samples). We identified 0-fold and 4-fold degenerate sites (here assumed as non-synonymous and synonymous loci respectively) by using the chromosome-level thrum assembly and its corresponding annotation with the script NewAnnotateRef.py (https://github.com/fabbyrob/science/tree/master/pileup_analyzers). Polymorphism was estimated separately for each gene at 0-fold non-synonymous and 4-fold degenerate synonymous sites using pixy. We compared π_*N*_, π_*S*_ and π_*N*_*/*π_*S*_ between *S*-locus-linked (LG10: 38,425,470-38,686,519) and neighboring genes (π_*N*_, π_*S*_: N=7 S-locus-linked genes and on each side of the S-locus; π_*N*_*/*π_*S*_: N=4 *S*-locus-linked and on each side of the *S*-locus after keeping only genes with π at 4-fold sites > 0) using Wilcoxon rank sum test (Supplementary Methods in Supplementary Note).

### Differential expression, gene set enrichment and patterns of expression of *S*-locus-linked genes

We analyzed differential expression between thrum and pin samples of floral buds (n thrum=6, pin=4), leaves (n thrum=6, pin=4), pistils (n thrum=3, pin=3), stamens (n thrum=3, pin=3) and petals (n thrum=3, pin=3) in the R package DESeq2 (Love et al. 2014), with logarithmic fold change correction using Approximate Posterior Estimation in the R package apeglm (Zhu et al. 2019) (Supplementary Methods in Supplementary Note). We controlled for FDR using the Benjamini-Hochberg method (Benjamini and Hochberg 1995) and considered genes with adjusted *P* ≤ 0.01 as significantly differentially expressed. Normalized counts of RNA mapped to *S*-locus-linked and DEG were compared between thrum and pin samples with a Wilcoxon rank-sum test. Gene set enrichment analyses were conducted in TopGO 2.46.0 using the weighted Fisher exact test (Alexa et al. 2006). We additionally investigated patterns of expression of *S*-locus-linked genes at mature floral organs by calculating their abundance as Transcript per Million (TPM), which allowed us to determine if transcripts from *S*-linked genes are expressed in pistils, stamens and petals. Gene expression was considered detected in a sample if TPM >= 0.5 percentile of TPM values for the sample, and genes were determined as expressed in the organ if present in two or more biological replicates. Finally, to better inform our understanding on the involvement of all genes clustered within the *S*-locus in controlling distyly, we leveraged the results of the functional annotation obtained with InterProscan and Blast (Supplementary Methods in Supplementary Note) to classify genes based on PANTHER family membership, and this information was used to manually retrieve GO terms using the AmiGO database (Ashburner et al. 2000; Carbon et al. 2009; Gene Ontology Consortium 2021).

### Identification of putative paralogs of S-linked genes and divergence estimates

We investigated the genomic distribution of the paralogs of *S*-linked genes by conducting Blast analyses (Altschul et al. 1990; Altschul et al. 1997; Camacho et al. 2009). Using the command ‘makeblastdb’, we created a custom data base including all protein-coding genes present in the annotation of the *L. tenue* assembly. Next, the longest isoforms of *S*-linked genes sequences were used as queries against the data base using the command ‘blastp’, and matches with an E-value =< 0.0 were deemed as significant. We kept the first ten best matches for each Blast result, and conducted reciprocal Blast analyses for each entry. Genes with the best matching results in both Blast and reciprocal Blast analyses were determined as the most likely paralogs of *S*-linked genes.

We estimated synonymous divergence between *S*-linked genes and their paralogs by extracting the coding sequence of the longest isoforms and aligned the sequences based on their translated amino acid sequences in webPRANK (Löytynoja and Goldman 2010). *S*-linked genes for which we could not reliably align their sequences with their corresponding putative paralogs were excluded from the analyses. We also excluded gene *LITEG00000052185* as the closest paralog was uncertain with nine highly similar paralogs detected across the genome (Table S8, Supplementary Table). The number of synonymous substitutions per synonymous site was estimated in MEGA X (Kumar et al. 2018; Stecher et al. 2020) using the Nei-Gojobori method (Nei and Gojobori 1986), excluding all alignment sites with gaps. Standard errors of synonymous divergence estimates were obtained based on 500 bootstrap replicates.

Finally, to investigate if homologs of *S*-linked genes are closely located or scattered throughout the genomes of other two species of the order Malpighiales, we used Phytozome’s Blast tool (https://phytozome-next.jgi.doe.gov/blast-search) to find the best matches of *L. tenue S*-linked genes in *Manihot esculenta* (v8.1) (Bredeson et al. 2016) and *Populus trichocarpa* (v4.1) (Tuskan et al. 2006). Finally, we conducted reciprocal Blast using the *L. tenue* protein data base for each of the matches previously obtained for both outgroup species.

## Supplementary Material

*Supplementary Note*

*Supplementary Tables*

## Acknowledgments

We thank Sara Mehrabi for assistance with lab work, José Ruíz-Martin for assistance with field work, and Jerker Eriksson for technical assistance. This project has received funding from the European Research Council (ERC) under the European Union’s Horizon 2020 research and innovation programme (grant agreement No 757451), from the Swedish Research Council (grant no. 2019-04452) to T.S., and from the Bergströms foundation to J.G.V. E.L.B. was funded through a Carl Tryggers grant to T.S. B.N. and V.E.K. are financially supported by the Knut and Alice Wallenberg Foundation as part of the National Bioinformatics Infrastructure Sweden at SciLifeLab. The authors acknowledge support from the National Genomics Infrastructure (NGI) in Stockholm and Uppsala (Uppsala Genome Center, SNP&SEQ), funded by the Knut and Alice Wallenberg foundation, the Swedish Research Council and Science for Life Laboratory. We acknowledge support of the Swedish National Infrastructure for Computing (SNIC) and Uppsala Multidisciplinary Center for Advanced Computational Science (UPPMAX) for assistance with massively parallel sequencing and access to computational infrastructure. Support by NBIS (National Bioinformatics Infrastructure Sweden) is gratefully acknowledged.

## Author Contributions

JGV, AL, AD, PWH, AF, BL, MA conducted experimental work.

JGV, MF, ELB, IB, LS, JD, AF, TS performed bioinformatic analyses.

OVP, BN, VEK, EP advised on sequencing and bioinformatic analyses.

JGV, AD, BL conducted field work.

JA, AB advised on study system and field work.

JGV and TS wrote the manuscript, with extensive input by MF and ELB. All coauthors commented on the manuscript.

TS designed the study and supervised the work.

