## Supplementary Note for "Genomic analyses of the *Linum* distyly supergene reveal convergent evolution at the molecular level"

### Content


**Supplementary Methods**

**Fig. S1.** Genome assembly and annotation flowchart

**Detailed methods description**

**Supplementary Results**

**Fig. S2.** Differences in coverage at the *S*-locus candidate region between morphs

**Fig. S3.** Alignments between haplotypes reconstructed for both morphs

**Fig. S4.** Genetic differentiation and differences in mean $\pi$ between morphs, population CL

**Fig. S5.** Recombination rates across LG

**Fig. S6.** Decay of linkage disequilibrium with distance at and around the *S*-locus

**Fig. S7.** GWAS results using FALCON-Unzip assembly

**Fig. S8.** Alignment between primary contig and haplotig harboring the candidate *S*-locus

**Fig. S9.** Manual editing of specific LG after anchoring step

**Fig. S10.** HiGlass manual edit

**Fig. S11.** Haplotypes and indel edge determination assisted by linked-read mapping

**Fig. S12.** Phased haplotypes of the ~260 kb hemizygous region

**Fig. S13.** Inferred formation of the gene set giving rise to the *Linum* distyly *S*-locus

**References**

### Methods

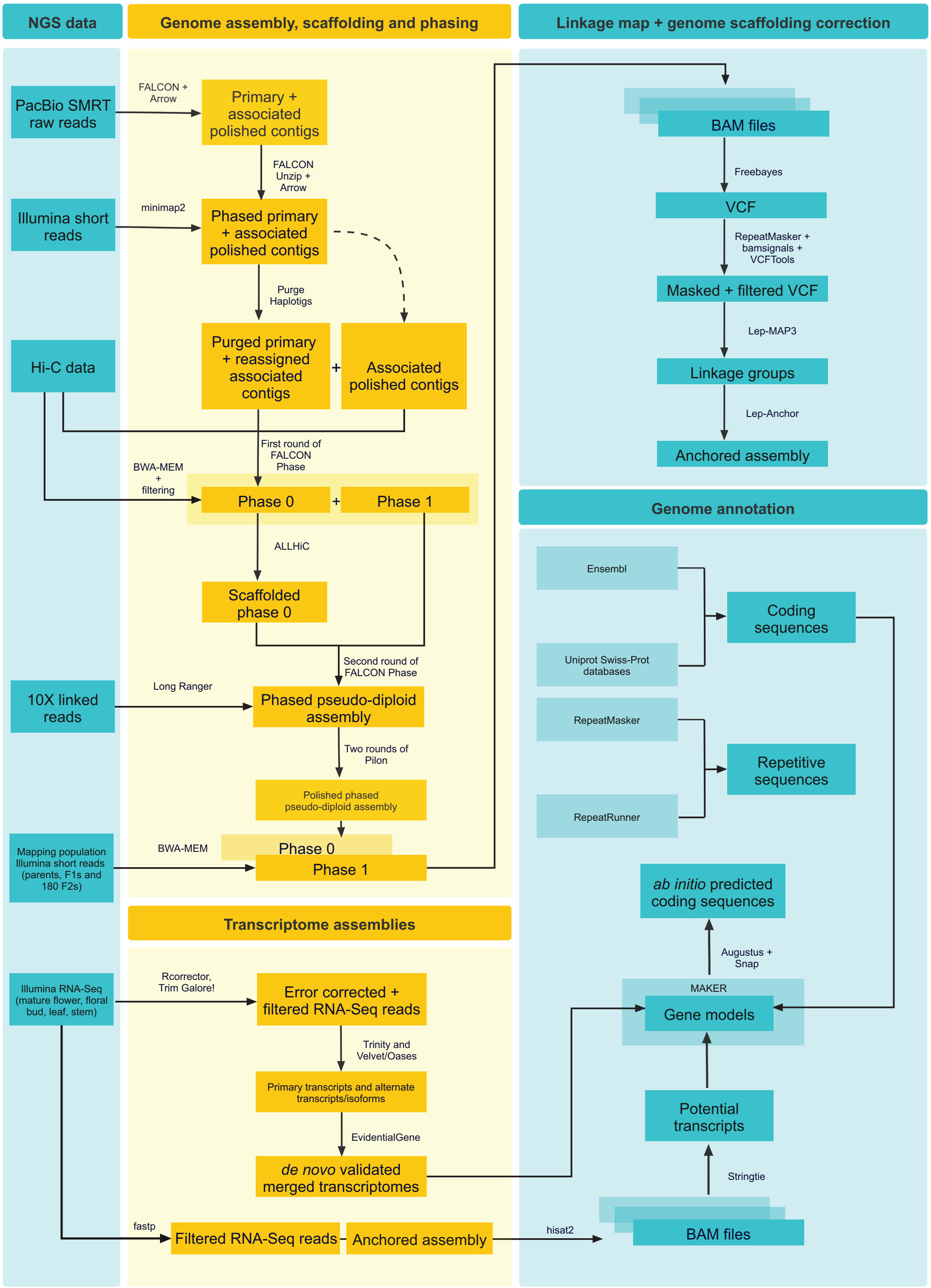


**Fig. S1. Genome assembly and annotation flowchart.** Flowchart illustrating the NGS data and bioinformatic pipeline used in the production of a chromosome-level genome assembly of *Linum tenue*.

### Supplementary Methods

### Detailed methods description

*Plant material*

Mature fruits of the distylous *Linum tenue* Desf. which occurs in southern Spain and north Africa were sampled at two localities in Andalusia, Spain: i) in Castillo de Locubín, Valdepeñas de Jaén (Jaén) (37°33'50.5"N 3°53'22.9"W) (CL onwards), and ii) Santa María de Trassierra, Sierra Morena (Córdoba) (37°55'55.1"N, 4°52'59.9"W) (SMT onwards). Fruits from each mother plant were collected in individual paper bags and treated with silica gel for 24 to 48 hours to reduce moisture. For plant propagation, we sterilized seeds using a treatment of 10% bleach solution with liquid detergent, followed by washes with 70% ethanol and sterile, distilled water. Seeds were germinated in Murashige-Skoog medium (Sigma Aldrich, USA) and covered with a thin layer of top agar. After 25 days of stratification at 4°C, we moved petri dishes to a controlled climate chamber at Stockholm University (Stockholm, Sweden) set to 16 h light at 20°C: 8 h dark at 18°C, 60% maximum humidity, 122 µE light intensity. We transplanted germinated seeds to pots containing a mixture of equal parts of vermiculite and perlite for two parts of regular soil (Hasselfors Garden, Sweden).

We collected and snap froze leaves of one thrum individual (SMT-34-3) for the production of a genome assembly, leaves of 43 individuals with known morph (CL: pin=8, thrum=8; SMT: pin=13, thrum=13) for population genomic analyses, and F1 (pin=F1.11 and thrum=F1.5) and 180 F2 individuals for the production of a linkage map (Table S7, Supplementary Tables). For generation of chromatin conformation and capture data, we sampled leaves from a thrum individual (individual SMT-24-4). These samples were processed as detailed in *DNA extraction and sequencing*.

For annotation of our genome assembly, we sampled and snap froze leaves, stems, floral buds and mature flowers of two thrum individuals (SMT-3-1 and SMT-38-2). Samples were taken with sterilized forceps and placed in 2 mL Eppendorf tubes, then flash frozen in liquid nitrogen. For differential expression analyses, we sampled and snap froze floral buds (*n* thrum=6, pin=4, three replicates per individual), leaves (*n* thrum=6, pin=4, three replicates per individual), pistils (*n* thrum=3, pin=3), stamens (*n* thrum=3, pin=3) and petals (*n* thrum=3, pin=3) (Table S7, Supplementary Tables). This material was used for various sequencing experiments detailed in *RNA extraction and sequencing*.

For a subset of three individuals, fresh leaves were collected and tested for absolute DNA content at Plant Cytometry Services (Didam, The Netherlands) using the propidium iodide (PI) staining method and flow cytometry using *Allium schoenoprasum* as standard (DNA content=15.03 pg/2C).

*Crosses and mapping population production*

We produced F1 plants by performing legitimate crosses between individuals from populations CL and SMT grown in our greenhouses at Stockholm University. After performing multiple hand-pollinations, we selected a single pair of parental plants (SMT.2.1 - Pin x CL.1.1 - Thrum) based on the number of successfully produced fruits harvested and stored as previously described. Two individuals from this F1 were used to produce an F2 mapping population. As before, we performed multiple crosses of a pair of F1 individuals (F1.11 - Pin x F1.5 - Thrum) and collected ripe fruits from plants that were grown and sampled following the same experimental settings and procedures previously described.

Finally, we grew 180 F2 offspring from this cross and sampled leaves for the production of a genetic map as detailed in the section *DNA extraction and sequencing*.

*DNA extraction and sequencing*

To generate a reference genome for the thrum morph of *L. tenue*, we produced a variety of data sets for which different extraction and sequencing methods were used. To obtain long-read sequencing data using the Single Molecule, Real-Time (SMRT) Sequencing technology from Pacific Biosciences (PacBio) (Sequel sequencing instrument, V3.0 chemistry), young leaves from a thrum individual (SMT-34-3) were snap frozen and disrupted by high-speed shaking with stainless steel beads. High molecular-weight (HMW) genomic DNA was extracted from approximately 100 mg of lysed material using the kit Genomic-tip 100/G (QIAGEN, Germany) following manufacturer's instructions. DNA concentration and purity were measured using Qubit and NanoDrop respectively. Sample integrity and fragment size distribution were determined through pulsed-field capillary electrophoresis using a Femto Pulse system. DNA was sheared to 60 Kb, and a SMRTbell library of size 31 Kb was generated and further sequenced on 16 SMRT cells. The resulting data was merged using DatasetMeger 0.3.0, and imported into the SMRT Analysis software suite (v2.3.0) (Pacific Biosciences, CA) to filter out sequences shorter than 500 bp or with a quality lower than 80, and generate subreads. This led to a data set of 119 Gb represented in 10,217,936 subreads (Table S7, Supplementary Tables).

To generate chromosomal conformation and capture data, nuclei from young leaves of a thrum individual were extracted using the protocol presented by Workman et al. (2018). Chromatin conformation capture data was generated following the manufacturer's instructions for the commercially available kit of the Hi-C method (Lieberman-Aiden et al. 2009) Proximo Hi-C 2.0 of Phase Genomics (Seattle, WA). First, intact nuclei were crosslinked using a formaldehyde solution, digested using the DpnII restriction enzyme, and proximity ligated with biotinylated nucleotides to create chimeric molecules composed of fragments from different regions of the genome that were physically proximal *in vivo*. Molecules were pulled down with streptavidin beads and processed into an Illumina-compatible sequencing library.

To polish the genome assembly, an HMW DNA extract from the individual SMT-34-3 (previously sequenced with SMRT PacBio) was used to create a 10X Chromium genomic library for linked-read sequencing. HMW DNA was extracted using the kit Genomic-tip 100/G (QIAGEN, Germany). Sequencing libraries were prepared from 0.6 ng of DNA using the Chromium Genome Library preparation kit (cat# 120~260/58/61/62) following the manufacturers' protocol (#CG00043 Chromium Genome Reagent Kit v2 User Guide). The library was sequenced on a HiSeqX system (paired-end 150bp read length, v2.5 sequencing chemistry). Linked-read sequences were also obtained for additional samples from natural populations (thrum=5 and pin=5) as previously explained. Both library preparation and sequencing were conducted at the SNP&SEQ Technology Platform in Uppsala (Sweden).

To aid scaffolding and correction of potential misassembles, we generated data for the production of a genetic linkage map. Thus, we sequenced the pair of parental plants (SMT.2.1 - Pin x CL.1.1 - Thrum), the F1 individuals (F1.11 - Pin x F1.5 - Thrum) and 180 F2 samples. Genomic DNA from young frozen leaves was extracted using the kit Isolate II Plant DNA (Bioline, UK). DNA concentration and purity were measured using Qubit and NanoDrop respectively. For parental individuals and F1s, sequencing libraries were prepared from 100 ng of DNA using the TruSeq Nano DNA sample preparation kit from Illumina (cat# 20015964/5) targeting an insert size of 350 bp, following the manufacturers' instructions. For F2s, indexed sequencing libraries were generated using the Nextera DNA Flex protocol, according to the manufacturers' instructions. For the sequencing of the parental individuals, libraries were sequenced on Illumina's HiSeqX sequencer (150-bp paired-end reads, v2.5 sequencing chemistry). For the sequencing of the F1 individuals, a NovaSeq 6000 system from Illumina was used with a S4 flowcell (150-bp paired-end reads, v1 sequencing chemistry). Finally, DNA from 180 F2 individuals was processed on a NovaSeq6000 (NovaSeq Control Software 1.6.0/RTA v3.4.4) sequencer with a 2x151 setup using 'NovaSeqXp' workflow in 'S4' mode flowcell. The Bcl to FastQ conversion was performed using bcl2fastq_v2.20.0.422 from the CASAVA software suite. For parental and F1 individuals, library preparation and sequencing were executed at the SNP&SEQ Technology Platform in Uppsala (Sweden) and for F2 individuals at the Genomics Applications Platform of the National Genomics Infrastructure (Sweden).

Finally, we generated a population genomic data set by sequencing 43 individuals with known morph (see *Plant material*; Table S7, Supplementary Tables). Genomic DNA extraction and short-read sequencing of these individuals was done as described above for parental plants.

*RNA extraction and sequencing*

For annotation of the genome assembly, total RNA was extracted from leaves, stems, floral buds and mature flowers using RNAeasy Plant Mini kit (QIAGEN, Germany) as per the manufacturer's instructions. RNA quantity and quality were evaluated by running aliquots of all samples on an Agilent Bioanalyzer 2100 (Agilent Technologies, Inc., Santa Clara, USA) using RNA Plant Nano microfluidic chips. Illumina TruSeq RNA Library v2 prep kits (Illumina, Inc., San Diego) were used for library construction. Two technical replicates derived from independently generated sequencing libraries were sequenced for each biological replicate (individual). Sequencing libraries were prepared using the TruSeq stranded mRNA library preparation kit (Illumina Inc.) including polyA selection according to the manufacturers' protocol. Sequencing was then performed on an Illumina NovaSeq S1 Sequencing System to produce paired-end 150bp read length sequences using the v1 chemistry at the SNP&SEQ Technology Platform in Uppsala (Sweden).

*Genome assembly*

The set of subreads that resulted from SMRT PacBio sequences were assembled using FALCON (Chin et al. 2016) and polished using the Arrow algorithm as implemented in Generate Highly Accurate Reference Contigs (GCpp, https://github.com/PacificBiosciences/gcpp) leading to the production of a preliminary assembly of 1,271 primary contigs (885.74 Mbp) and 597 associated contigs (90.25 Mbp) (Fig. S1 and Supplementary Methods, Supplementary Note; Table S1, Supplementary Tables).

The FALCON assembly was phased with FALCON Unzip (combo package pb-falcon v0.2.4, https://github.com/PacificBiosciences/pbbioconda) (Chin et al. 2016) and polished using the SMRT PacBio subreads with Arrow as implemented in Generate Highly Accurate Reference Contigs (GCpp, https://github.com/PacificBiosciences/gcpp) leading to the production of a primary assembly of 879.49 Mbp (No. sequences=1,222) and a set of haplotigs of 161.13 Mbp (No. sequences=1540) (Table S1, Supplementary Tables).

*Haplotig purging*

Diploid-aware assemblers such as FALCON-Unzip can fail to identify homologous regions when local heterozygosity is very high. In other words, haplotigs that should be assigned to a certain primary contig are misclassified as a different region in the genome. We used the pipeline Purge Haplotigs (Roach et al. 2018) to reassign allelic contigs based on read-depth analyses and repeat annotations. Illumina short-reads obtained from individual SMT-34-3 were mapped to the FALCON-Unzip reference genome containing only primary contigs using minimap2 (Li 2018), and the resulting alignments were processed with the function purge_haplotigs to generate a coverage histogram for contigs in the assembly. The resulting bimodal histogram allowed to determine the read-depth cutoff values (low=15, mid=45 and high=85) used to flag contigs that could be potentially removed using the function purge_haplotigs cov. Finally, the output coverage stats files were processed with purge_haplotigs purge to perform contig reassignment, and the handling of repetitive regions was facilitated using the library of repeats of *L. usitatissimum* available in RepeatMasker (4.1.0) (Smit et al. 2013). The purging of the primary assembly led to the reassignment of 259.47 Mbp of the genome sequence to the set of haplotigs (Table S1, Supplementary Tables). Contig 000228Flarrow, preliminarily identified as a candidate for the distyly *S*-locus (see *Preliminary Identification of an S-locus candidate region* below), was determined as an haplotig of 000175Flarrow, suggesting that these contigs harbored highly divergent yet homologous regions (Fig. S8 in Supplementary Results).

*Scaffolding and phasing of the genome assembly using Hi-C data*

The purged version of the primary assembly (615.74 Mbp, No. sequences=553), the set of haplotigs originally identified with FALCON-Unzip concatenated to those reassigned by PurgeHaplotigs (420.59 Mbp, No. sequences=2128), and the Hi-C data were used to phase and scaffold the genome assembly. In the first round of phasing, the partially phased PacBio-based assembly was processed together with the Hi-C data using FALCON-Phase v1.2.0 (Kronenberg et al. 2021) available as the pb-assembly package (v0.0.8) from Bioconda (https://bioconda.github.io/recipes/pb-assembly/) to produce two full length pseudo-haplotypes for phase0 and 1. For scaffolding, Hi-C reads were aligned to phase0 of the assembly using BWA-MEM (v0.7.17) (Li and Durbin 2009) followed by filtering with samtools (v1.9) (Li et al. 2009) and the script PreprocessSAMs.pl (https://github.com/tangerzhang/ALLHiC/blob/master/scripts/) to keep only links with strong signals in the Hi-C data set. Then the scaffolds of phase0 were chained into ten pseudochromosomes using ALLHiC (v0.9.13) (Zhang et al. 2019) following these four steps: partition, rescue, optimize and build. After a second round of FALCON-Phase using the scaffolded version of phase0 as primary assembly and the non-scaffolded version of phase1 as associated contigs, misplaced scaffolds were reassigned to each phase. This yielded a chromosome-scale phased pseudo-diploid assembly of ten pseudochromosomes totaling a sequence of 703.83 Mb for phase0 and 702.16 Mb for phase1 (Fig. S1 and Supplementary Methods; Table S1, Supplementary Tables). Finally, to polish the assembly, 10X linked reads were aligned to each phase of the reference using Long Ranger (https://github.com/10XGenomics/longranger), followed by two rounds of Pilon (Walker et al. 2014). Assembly statistics for each step of the assembly were obtained using the script gaas_fasta_statistics.pl (https://github.com/NBISweden/GAAS/blob/master/bin/).

*Preliminary identification of an S-locus candidate region*

The production of a phased-diploid assembly can be especially difficult in highly heterozygous regions. We used a partially-phased genome assembly to conduct preliminary identification of *S*-locus candidate regions, to ensure that relevant candidate regions were retained in subsequent steps towards the production of a phased chromosome-level assembly. For this purpose, we conducted a preliminary genome-wide association analysis (GWAS) of whole-genome short-read DNA sequences from 43 individuals with known morph (see *Plant material* and *DNA extraction and sequencing*).

Short read adapter removal and quality trimming steps were conducted using `bbduk` from BBMap/BBTools (Bushnell 2015). Trimmed paired-end reads were mapped to the polished version of the FALCON assembly with BWA-MEM (v0.7.17) skipping alignments with mapping quality lower than 20 (Li and Durbin 2009), and duplicated reads were removed using `MarkDuplicates` from Picard tools v2.0.1 (Broad Institute 2019).

BAM files were processed with samtools/bcftools (Danecek and McCarthy 2017) to produce genotype likelihoods from sequence alignments using the function `mpileup`, and the VCF file containing variants (SNPs/INDELs) and invariant sites was created with the function `call`. The resulting VCF file was further processed to only keep biallelic SNPs and invariant sites, and filtered based on quality (QUAL > 20), coverage (AVG(FMT/DP) > 10 & AVG(FMT/DP) < 50) and data missingness (F_MISSING < 0.2) using the function `filter` from bcftools. Last, repetitive regions were identified using the library of *L. usitatissimum* available in RepeatMasker (4.1.0) (Smit et al. 2013) and removed from the VCF using the function intersect from BEDTools.

We used the resulting VCF file to test for an association between morph and SNP genotype in PLINK (v1.90b4.9) ( www.cog-genomics.org/plink/1.9/, Chang et al. 2015) using Fisher's exact test, with significance adjustment using the Benjamini-Hochberg procedure to control the False Discovery Rate (FDR). This analysis led to the identification of 299 SNPs significantly associated with floral morph ($P < 0.01$), of which 99.67% were mapped to contig 000228F_arrow (502,875-875,960 bp) (Fig. S7). Additionally, the alignment between the DNA sequences of 000228Flarrow, identified as an haplotig of 000175Flarrow in the haplotig purging step (see *Haplotig Purging* above and Fig. S8), indicated the existence of a ~260 kb region in 000228Flarrow (614,903-872,429) that was absent in contig 000175Flarrow and that showed strongly significant association with morph (Fig. S8). Given this result, we proceeded to keep sites linked to 000228Flarrow in further steps of the assembly of the genome. To do this, it was necessary to manually edit the final assembly (after scaffolding correction), as the first round of FALCON-Phase resulted in the removal of this region from the assembly, possibly because it was classified as secondary and discarded.

*S-locus location and haplotype phasing*

After scaffolding correction based on the linkage map (see section *Linkage map production and scaffolding correction)* we mapped the contig 000228Flarrow against the reference genome using minimap2 (Li 2018) and inserted the ~260 kbp region in its corresponding location at LG10:38,426,515-38,684,012. To check this edit and phase haplotypes in this region we mapped 10X linked reads from individual SMT-34-3 to the reference genome using Long Ranger (https://github.com/10XGenomics/longranger) which through freebayes (Garrison and Marth 2012) calls the phased variants. The ~260 kb hemizygous region is contained in a 1 Mbp phased block (LG10:37,829,718-38,865,685). We visually inspected the mapped 10X linked-reads against the ~260 kb indel with Loupe (https://support.10xgenomics.com/genome-exome/software/visualization/latest/what-is-loupe). Haplotypes and edges of the ~260 kb indel were identified upon visual inspection of linked-reads mapped to this region (Fig. S11 in Supplementary Results). To ensure that the reference genome carries the dominant *S*-locus haplotype, we modified it based on phased SNPs and small indels called by FreeBayes that belong to the phased haplotype of interest (Fig. S12 in Supplementary Results).

*10x Genomics linked-reads Supernova assemblies*

We *de novo* assembled genomes for samples of both morphs (thrum=4, pin=5) with the Supernova pipeline (https://github.com/10XGenomics/supernova) using the default parameters to obtain the output type pseudohap2; the summary statistics of the assembly are in the (Table S10, Supplementary Tables). We mapped the region LG10: 38,180,000-38,930,000 against each 10x Genomics linked-reads Supernova assembly to identify the ~260 kb hemizygous region using minimap2 (Li 2018).

*Linkage map production and scaffolding correction*

Short Illumina reads from individuals used for the production of the mapping population (180 F2, the F1s F1.11 x F1.5, and the parents SMT.2.1 x CL.1.1) were aligned to the primary assembly using BWA-MEM with default parameters and the resulting sam files were converted to bam format using samtools. Read groups were added, the files were sorted, and duplicates were removed using Picard tools v2.0.1 (Broad Institute 2019). Variants were called using FreeBayes (v.1.3.2) specifying a minimum coverage of 5 reads and the use allele mapping quality when calculating data likelihoods (Garrison and Marth 2012).

The resulting VCF file was filtered in successive steps. To remove repetitive regions that might be collapsed in the genome assembly, a library of repetitive regions was created for the latest version of the assembly using the NCBI BLASTDB database implemented in the repeat modeling pipeline RepeatModeler (v1.0.11) (Smit and Hubley 2008). The identified repetitive regions were masked in the assembly with the option for slow search (-s) from RepeatMasker (Smit et al. 2013), and further removed from the VCF. Then, we removed SNPs and invariant sites in windows with unusually high coverage. For each sample, read depth was calculated in 500 bp windows using the R package bamsignals (v1.26.0) (Mammana and Helmuth 2021), and windows with a coverage 2x higher than the within-sample median in at least in half of all individuals were removed. Further filtering for phred score (>20), the ratio of phred score over count of observations of the alternate haplotype (>10), the ratio of reference alleles to total alleles in heterokaryotypes (<0.25, >0.75), the number of alternate alleles noted on the forward and reverse strands (>0), the number of reads placed right and left (>1), and the ratio of mapping quality of alternate over reference alleles (>0.9, <1.05) was done using vcflib (Garrison et al. 2021). As we were only interested in segregating sites, VCFtools (Danecek et al. 2011) was used to remove sites with a minor allele frequency < 0.2. We also removed indels, sites with more than 2 alleles and sites with more than 20 missing genotypes (~10%). Finally, sites were thinned out in VCFTools (Danecek et al. 2011) so that all SNPs were at least 1 kb apart (--maf 0.2 --max-missing-count 20 --max-alleles 2 --remove-indels --thin 1000).

DNA extracts from four of the F2 individuals were sequenced twice in the same batch and used to identify and retain sites where all four pairs had present and matching genotypes based on these four duplicated individuals. Additional non-informative sites were removed after identifying sites where parents and F1s sequences had non-matching genotypes based on heterozygous markers. Segregation distortion was estimated for each marker using a chi-squared test, p-values were adjusted using the Benjamini-Hochberg method (1995) for FDR as implemented in the R function p.adjust(), and taking a stringent approach, we only kept sites associated with p-values > 0.1. These filtering steps resulted in a final set of 5,857 SNPs used for linkage map construction.

The generation of linkage groups relied on Lep-MAP3 (Rastas 2017) using a Logarithm of Odds (LOD) limit of 30 and a theta value of 0.5. We were able to place 5,826 markers into 10 linkage groups. Before moving to Lep-Anchor we generated a map to check the distribution of markers across the linkage groups. We ran the marker ordering for all linkage groups 10 times and retained the order with the lowest log likelihood. We then checked our map and removed markers if there were less than 5 markers from one of our scaffolds in a linkage group. This step resulted in the removal of 33 markers, for and thus the final number of markers used was 5,793. We remade our linkage groups, then input our data into Lep-Anchor (Rastas 2020) along with a chains file made in HaploMerger (Huang et al. 2012) using the masked reference and a paf file made by mapping the first version of the FALCON phased version of the genome against the reference with minimap2 (Li 2018) in order to assign primary scaffolds to pseudochromosomes and correct misjoins.

We inspected the results from Lep-Anchor using Marey maps and identified three regions, (LG1:10,941,743-18,235,911, LG5:1-2929725, and LG3:12,307,570-20,979,995) where the recombination distance and the physical position did not match (Fig. S9 in Supplementary Results). We used this information to manually edit the assembly for LG1 and LG2. In the first case we moved this region to the end of LG1 and in the second we inserted it at position 56,376,345 of LG5. This editing was supported by our Hi-C data based on a HiGlass visualization (Kerpedjiev et al. 2018) (Fig. S10 in Supplementary Results). In the case of LG3, the discrepancy between recombination map and physical position showed a complex pattern that was not feasible to correct manually with the currently available data. The summary statistics of the final version of the chromosome-level assembly (i.e. genome fragmentation, FASTA file checking, BUSCO scores, GC content, etc.) were obtained using the Nextflow pipeline AnnotationPreprocessing from the github repository https://github.com/NBISweden/pipelines-nextflow with the parameter file params.config.

*Organelle genomes*

To assemble chloroplast and mitochondrial genomes we used 10X linked-read data from individual SMT-34-3 and the software GetOrganelle (Jin et al. 2020).

*Genome annotation*

Prior to the annotation of the assembly sequence, genome completeness was assessed with BUSCO (v4.0.2) (Waterhouse et al. 2018) using the eudicots_odb10 gene dataset, and our estimate indicated that most of the expected genes (94.2%) are present in the assembled genome sequence. The identification of coding sequences relied on the usage of curated and custom protein sequences databases, and the assembly of the transcriptomes from different tissues. We used the Uniprot Swiss-Prot database (Magrane and Consortium 2011) to produce a non-redundant protein sequence database that consisted of 563,972 manually curated proteins (downloaded on December of 2020). We also identified and downloaded taxon-specific protein data bases from UniProt and Ensembl, including the proteomes of Rosids (rosids_swissprot.fasta with 22,046 proteins, and Vitis_vinifera.12X.pep.all.fa, with 29,927 proteins), the order Malpighiales (Populus_trichocarpa.Pop_tri_v3.pep.all.fa with 73,012 proteins) and the genus *Linum* (Lusitatissimum_200_v1.0.protein.fa with 43,484 proteins).

In addition to protein-based evidence, we generated *de novo* transcriptome assemblies for annotation using RNA-Seq data from thrum mature flowers, floral buds, leaves and stems (see *RNA extraction and sequencing*). Raw reads were processed with Rcorrector (Song and Florea 2015) using default settings to correct errors in the reads, and reads containing non-correctable errors were dropped from the dataset. Adapter and quality trimming were then conducted using Trim Galore! (v0.4.4) (https://github.com/FelixKrueger/TrimGalore) (settings: paired mode, retain unpaired reads, minimum sequence length=36, trimming ends of reads below quality=5). Ribosomal reads were removed from the dataset through alignment to a custom database comprising the SILVA LSU, SSU (Pruesse et al. 2007; Quast et al. 2013) and 5SRNAdb (Szymanski et al. 2016) ribosomal databases. Alignments for filtration were performed with Bowtie2 (v2.3.3.1) (Langmead and Salzberg 2012) using the option `--very-sensitive-local`. All aligned reads (i.e. retrieved as ribosomal) were discarded, while the unaligned ones were kept for *de novo* assembly with Trinity (2.9.1) using the default settings (Grabherr et al. 2011; Haas et al. 2013). Additional assemblies were produced by processing the *in silico* normalized files created through Trinity with Velvet (Zerbino and Birney 2008) and Oases (Schulz et al. 2012) using as *k*-mer values all odd numbers in the range between 21-49. The transcriptomes obtained with the pipelines of Trinity and Velvet/ Oases were merged using EvidentialGene to identify the primary transcripts and alternate transcripts/isoforms accepted as valid through the merging pipeline.

We generated transcriptomes for mature flowers, floral buds, leaves and stems from thrum individuals (see *RNA extraction and sequencing*) using a reference guided assembly approach. RNA-Seq data and the chromosome-level reference genome were processed through the pipeline TranscriptAssembly (https://github.com/NBISweden/pipelines-nextflow/tree/master) that uses fastp (Chen et al. 2018) for quality control and preprocessing of raw files, hisat2 for the alignment of RNA reads (Kim et al. 2015), and StringTie (Pertea et al. 2015) to assembly transcripts from each tissue independently.

To avoid issues stemming for the overlapping of repetitive sequences and genes, and to avoid non-specific gene hits, we created a custom repeat library modelled using RepeatModeler (Smit and Hubley 2008) and RepeatMasker (Smit et al. 2013) as previously explained. Since protein-coding genes can contain repetitive sequences, the library of repeats was vetted against the protein set (after transposons' removal) to exclude any nucleotide motif present in low-complexity coding sequences. The final identification of repetitive sequences in the genome was conducted using RepeatMasker (Smit et al. 2013) and RepeatRunner (Smith et al. 2007), allowing the identification of highly divergent repeats and protein coding portions.

Gene models were constructed using MAKER (Holt and Yandell 2011) guided with evidence from both aligned transcript sequences and reference proteins, and then were used to train the *ab inito* prediction tools (https://github.com/NBISweden/pipelines-nextflow/tree/master/AbinitioTraining). Once the *ab initio* tools were trained, a new run of MAKER was conducted. Two approaches were used: first, the combination of the *ab initio* tools Augustus (v3.4.0) (Stanke et al. 2008) and Snap (Korf 2004), and second, a prediction using only Augustus. These strategies were compared based on the predicted number of genes and the visual inspection of the annotated features to examine the occurrence of false positive predictions, and the presence of fragmented and missing genes. The approach using only Augustus lead to a higher percentage of genes identified compared with the Augustus + Snap gene build (93.8 vs 91.6%, respectively) (Table S2, Supplementary Tables), and the visual inspection of the results showed a clear reduction in false positive prediction. Hence, the Augustus only-based was retained for further analyses. Gene model quality-control was estimated as the Annotation Edit Distance (AED) (0= perfect agreement of the annotation to aligned evidence, 1= no evidence supporting the annotation) with MAKER (Holt and Yandell 2011), and this information was included for each retrieved mRNA within the genome annotation files (.gff).

Functional annotation of the translated CDS features of was conducted using the pipeline FunctionalAnnotation (https://github.com/NBISweden/pipelines-nextflow/tree/master/FunctionalAnnotation). This pipeline uses Blast and InterProScan (Hunter et al. 2012) to retrieve information on protein function from 20 different sources, which was associated to each mRNA feature. To infer gene and protein names, protein sequences were blasted against the Uniprot/Swissprot reference data set, and hits with the best score (E-values < $1x10^{-6}$) were kept. The annotation of tRNA sequences relied on tRNAscan (1.3.1) (Lowe and Eddy 1997) followed by the removal of features with AED scores equal to 1. Finally, other ncRNAs were predicted using the database Rfam (Nawrocki et al. 2015) using only highly conserved eukaryotic ncRNA families, and the co-variance models provided by Rfam were then processed using Infernal (Nawrocki and Eddy 2013).

We performed a contamination analysis of the reference genome using Kraken2 (Wood et al. 2019). The analysis was conducted in 10 kb windows using the UniVec_Core database. We detected a region contaminated with thrip sequences (LG3:75,714,800-75,822,800) and we hard-masked it with BEDTools (Quinlan and Hall 2010).

*Manual annotation of S-locus candidate region*

Prior to the annotation of the reference genome we determined that the sequence of TSS1, earlier reported as a thrum-specific protein in *L. grandiflorum* (Wood et al. 2019), mapped to the ~260 kb-region linked to the candidate distylous *S*-locus (significant alignment with NCBI BLASTP: E-value=$3x10^{-6}$, sequence identity=43%, length=173, gaps=8%). The sequence of *L. grandiflorum TSS1* was downloaded from GenBank (accession code: AB617824.1, https://www.ncbi.nlm.nih.gov/) and blasted against the chromosome-level thrum assembly with blast (v2.7.1+) (Altschul et al. 1990) using the algorithm megablast with soft masking using a masking data base created with dustmasker (Zhang et al. 2000). Despite this result, *TSS1* was not retrieved by the automatic pipeline, suggesting that this and other genes linked to the distyly *S*-locus might be absent in the annotation. Thus, we conducted manual annotation of the region LG10:38,426,515-38,684,012. First, genes were predicted in the sequence spanning LG10 between 38.40 and 38.70 Mbp with Augustus (Stanke et al. 2008) using the training conducted in *Arabidopsis thaliana* and the option `--softmasking=0`. Then, we sought additional evidence supporting the existence of the predicted protein coding sequences by using RNA sequencing data from leaves, stems, floral buds and mature flowers from thrum individuals (two biological replicates sequenced in duplicate). The CDS and exon sequences for each gene feature obtained with Augustus were processed together with the thrum chromosome-level genome assembly using STAR to generate the genome index (Dobin et al. 2013), and RNA sequences from each sample were independently processed to count the number of reads mapped to each gene by setting `--outFilterMultimapNmax=1` to exclude multi-mapping reads. Files listing the number of reads mapped to gene features (ReadsPerGene.out.tab) were processed to identify coding sequences that were supported as present in the transcript of at least one tissue (i.e., if present in two or more of the replicates). Since genes present in the hemizygous region (LG10: 38,426,515-38,684,012) should only get thrum-derived sequences mapping to it, we repeated these analyses using pin-derived sequences to identify and remove false annotations. This procedure resulted in the removal of one gene annotation. If a certain protein coding sequence was present both in the automatic and manual annotations, we kept only one of them for downstream analyses. Finally, the functional annotation of these protein coding genes was conducted as described in *Genome annotation*.

*Identification of regions showing coverage differences between thrum and pin sequences*

Whole-genome sequences (Illumina short reads) from 43 individuals with known morph (CL samples: pin=8, thrum=9; SMT samples: pin=13, thrum=13) were quality and adaptor trimmed and mapped to the scaffolded and phased assembly as described in *Preliminary identification of an* S*-locus candidate region.* Per-site coverage was estimated with BEDTools (Quinlan and Hall 2010) for each BAM file, and sites in repetitive regions were identified with the repeats library from *L. usitatissimum* from RepeatMasker (Smit et al. 2013) were removed. We used the library of repeats from homostylous *L. usitatissimum* instead of the repeats from the *L. tenue* assembly to avoid the removal of regions that might be linked to the distyly *S*-locus, which could be enriched in repetitive sequences. We divided the repeat-masked version of the genome assembly into 50 kb windows, and estimate the mean, the median and the sum of coverage per window for each sample independently suing the function `map` from BEDTools (Quinlan and Hall 2010). The resulting per-window summarized estimates of coverage were processed to estimate normalized mean and median coverage using genome-wide estimates of mean and median coverage per sample, respectively. Additionally, we identified the LG showing the highest number of windows showing elevated thrum: pin normalized mean and median coverage (>1.5X), and produced plots that were used to visually detect adjacent windows in which thrum samples showed consistently higher normalized mean and median coverage than pin samples. Finally, we tested for significant differences in normalized mean and median coverage between thrum and pin samples for each 50 kb window using a two-sided Wilcoxon signed-rank test followed by Bonferroni correction.

*Estimates of genetic divergence and differentiation between thrum and pin-derived sequences*

BAM files produced for the coverage analyses were further processed to generate a VCF file containing both biallelic SNPs and invariant sites, followed by filtering based on quality (QUAL > 20), coverage (AVG(FMT/DP) > 10 & AVG(FMT/DP) < 50) and data missingness (F_MISSING < 0.2) sites as explained in the section *Preliminary identification of an* S*-locus candidate region*. The identification of repetitive regions to be masked in further analyses relied on two approaches. First, we created a custom library of repeats from the chromosome-level thrum *L. tenue* assembly using the NCBI BLASTDB database (consensi.fa.classified) as implemented in RepeatModeler (Smit and Hubley 2008), and these repetitive regions were masked using the slow search option (-s) with RepeatMasker (Smit et al. 2013) to create a BED file listing the LG, start and ending position of repetitive sequences. Second, using the same approach described in *Identification of regions showing coverage differences between thrum and pin sequences*, we identified 10 and 50 kb windows showing normalized mean coverage values higher than four (i.e. local coverage four times higher than genome-wide coverage estimates), which likely represent highly repetitive regions that were not identified using the first strategy. Since the presence of repeats in the candidate distyly *S*-locus might result in mapping ambiguities that inflate local polymorphism estimates, we masked the ~260 kb (~260,497 bp) hemizygous region previously identified in thrum individuals (LG10: 38,426,515- 38,68,4012) (see *Identification of regions showing coverage differences between thrum and pin sequences*, Supplementary Methods). The genomic coordinates of the hemizygous region were further confirmed after repeating the coverage analyses using 100 bp windows, which aided the identification of loci in which the thrum: pin normalized mean coverage drastically increased. The repeat-masked resulting VCF file was processed with pixy (Korunes and Samuk 2021) to estimate $F_{ST}$ and $d_{xy}$ (between thrum and pin samples), and $\pi$ (to estimate the difference in nucleotide diversity between thrum and pin samples) in 5 Kb windows. Each population (CL=17 and SMT=26 samples) was analyzed independently, and sample-morph association files were provided to pixy in the argument `--populations`. To identify which windows show significant $F_{ST}$ and differential $\pi$ between morphs, we conducted permutation tests. Approximate test-statistic distributions for each window were obtained by iterating the list of sample ID – morph association 1000 times, and the observed value was compared with the distributions to calculate the P-values, followed by a correction for multiple testing with the FDR method using the Benjamini-Hochberg approach.

*GWAS*

We conducted a GWAS to investigate whether loci flanking the ~260 kb hemizygous region are associated to floral morph. Thus, we used only biallelic SNPs derived from the VCF file generated for the analyses described in *Estimates of genetic divergence and differentiation between thrum and pin-derived sequences* for this analysis. A data set including 42 individuals from CL and SMT (equal number of samples per morph in each population) was processed as described in *Preliminary identification of an* S*-locus candidate region*. After correction for multiple testing with the method FDR using the Benjamini-Hochberg procedure (Benjamini and Hochberg 1995), Manhattan plots depicting genomic position *vs* -$\log_{10}$(*P*-value) were produced for all LG together, LG10 and the candidate region based on the coordinates identified as significantly associated to morph around the ~260 kb hemizygous region.

*Comparison of* $\pi_N/\pi_S$ *between distyly* S*-locus-linked and neighboring genes*

To investigate if $\pi_N/\pi_S$ estimates differ between distyly *S*-locus-linked and neighboring genes, we produced a thrum-only VCF file containing biallelic SNPs and invariant sites (CL=9 and SMT=13 samples), which was obtained using the same mapping, variant calling and preliminary filtering steps described in *Estimates of genetic divergence and differentiation between thrum and pin-derived sequences*. To avoid the removal of loci in coding sequences that might overlap with repetitive regions, the list of repeats in identified with RepeatModeler (Smit and Hubley 2008) and RepeatMasker (Smit et al. 2013), and the detection of 10 and 50 Kb windows with normalized mean coverage values higher than four (see *Estimates of genetic divergence and differentiation between thrum and pin-derived sequences*) was intersected with the coordinates of all annotated genes (see *Genome annotation*) using the option -v (only report ranges in A that does not overlap with ranges in B) from BEDTools (Quinlan and Hall 2010), and the resulting file was used to mask the VCF file.

We identified 0-fold and 4-fold degenerate sites (here assumed as non-synonymous and synonymous loci respectively) by using the chromosome-level thrum assembly and its corresponding annotation with the script NewAnnotateRef.py (https://github.com/fabbyrob/science/tree/master/pileup_analyzers). Then, the repeats-masked VCF was processed with pixy (Korunes and Samuk 2021) to estimate $\pi$ for each gene reported in the annotation by restricting the analyses to sites listed as 0-fold and 4-fold sites, independently. The resulting files were processed to compare $\pi_N$, $\pi_S$ and $\pi_N/\pi_S$ between *S*-locus-linked and neighboring genes using a Kruskal-Wallis rank sum test. We defined as *S*-locus-linked genes those annotated in the region spanning LG10: 38,425,470-38,686,519, i.e. ~260 kb of the hemizygous region and morph associated-loci flanking the indel based on GWAS (*n*=7 genes with 4-fold > 0). Neighboring genes correspond to those adjacent to the *S*-locus both in down- and upstream regions (*n*=7 genes with 4-fold > 0 on each side of the *S*-locus).

*Comparison of TE enrichment between the distyly* S*-locus and neighboring windows*

The chromosome-level genome assembly and its corresponding annotation of repetitive elements (see *Genome annotation*) were processed to estimate and compare the percentage of TEs between S-locus-linked and neighboring loci (as defined in *Comparison of* $\pi_N/\pi_S$ *between distyly* S*-locus-linked and neighboring genes*). Microsatellites (STRs and simple repeats) and regions of low complexity were removed from the analyses, and the fraction of TEs in 25 kb windows were compared between the *S*-locus and the neighboring regions using a Wilcoxon rank-sum test. Finally, the annotation of repetitive elements was processed to characterize the most abundant types of TEs (DNA transposons, LINE, LTR, RC or other) for all LG, LG10 and the *S*-locus.

*Differential expression analyses*

RNA sequencing datasets from floral buds (*n* thrum=6, pin=4, three replicates per individual), leaves (*n* thrum=6, pin=4, three replicates per individual), pistils (*n* thrum=3, pin=3), stamens (*n* thrum=3, pin=3) and petals (*n* thrum=3, pin=3) were obtained using the same protocols described in *RNA extraction and sequencing*. Adapters removal and QC trimming steps were conducted using `bbduk` from BBMap/BBTools (Bushnell 2015), and the resulting files were processed as described in *Manual annotation of S-locus candidate region* to obtain counts on the number of reads mapped to gene features (ReadsPerGene.out.tab). To determine if replicates obtained from buds and leaves showed similarity between them, we estimated the Spearman's rank correlation between counts for sequences derived from the same individual – tissue combination in R (R Core Team 2021).

In order to conduct differential expression analyses, the ReadsPerGene.out.tab files were processed in R to identify DEG between pin and thrum samples for each tissue using the R package DESeq2 (Love et al. 2014). Since the sequencing experiment for floral buds and leaves was conducted in triplicate, counts coming from the same tissue and individual were added up prior to DE estimation. Results from the DE analyses were further processed to more accurately estimate logarithmic fold change values using the method Approximate Posterior Estimation for generalized linear model implemented in the R package apeglm (Zhu et al. 2019). Genes with estimates of a $P$ value < 0.01 (after adjustment with the FDR/Benjamini-Hochberg method) were deemed as differentially expressed. Finally, we obtained normalized counts for *S*-locus-linked genes for each tissue and compared between morphs using a Wilcoxon rank-sum test.

*Detection of* S*-locus-linked genes expression in mature floral tissues*

Files with transcripts counts, previously obtained for the differential expression analyses (see *Differential expression analyses*), were used to estimate and compare the expression level of *S*-locus-linked genes in each sample. Transcript per Million (TPM) values were calculated in R. The annotation was filtered using the function agat_sp_keep_longest_isoform.pl from AGAT (Dainat et al. 2022) to identify the longest isoform per gene and use this information to control for gene length. Genes were deemed as expressed within samples if TPM values were higher than the 0.5 percentile of the distribution of TMP values of all genes for that sample. Finally, transcripts were identified as present in a certain tissue if they were present in two or more biological replicates.

### Results

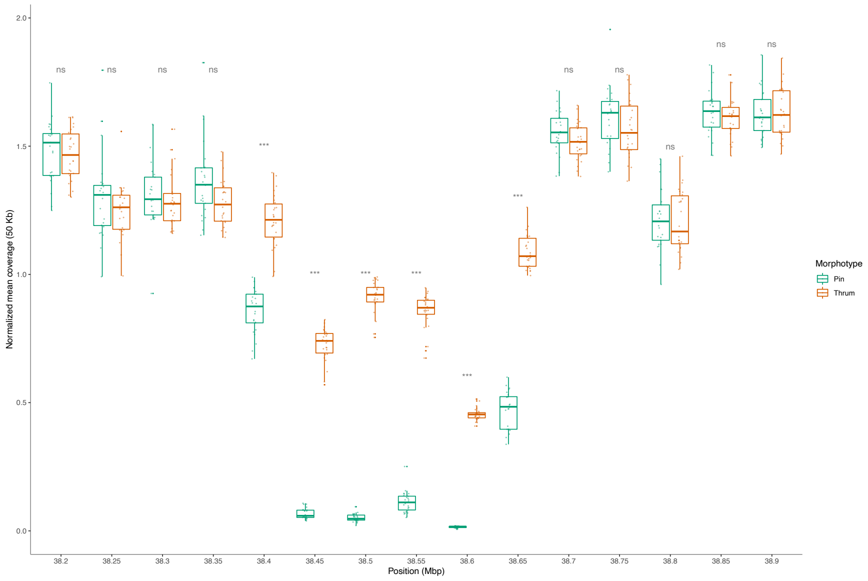


**Fig. S2. Differences in coverage at the S-locus candidate region between morphs.** Comparison of normalized mean coverage between thrum ($n$=26) and pin ($n$=25) samples across 15 windows (50 Kb) spanning the S-locus and 250 Kb downstream and upstream regions (Wilcoxon test, ns: nonsignificant, *: $P$<0.05, **: $P$<0.01, ***: $P$<0.001).

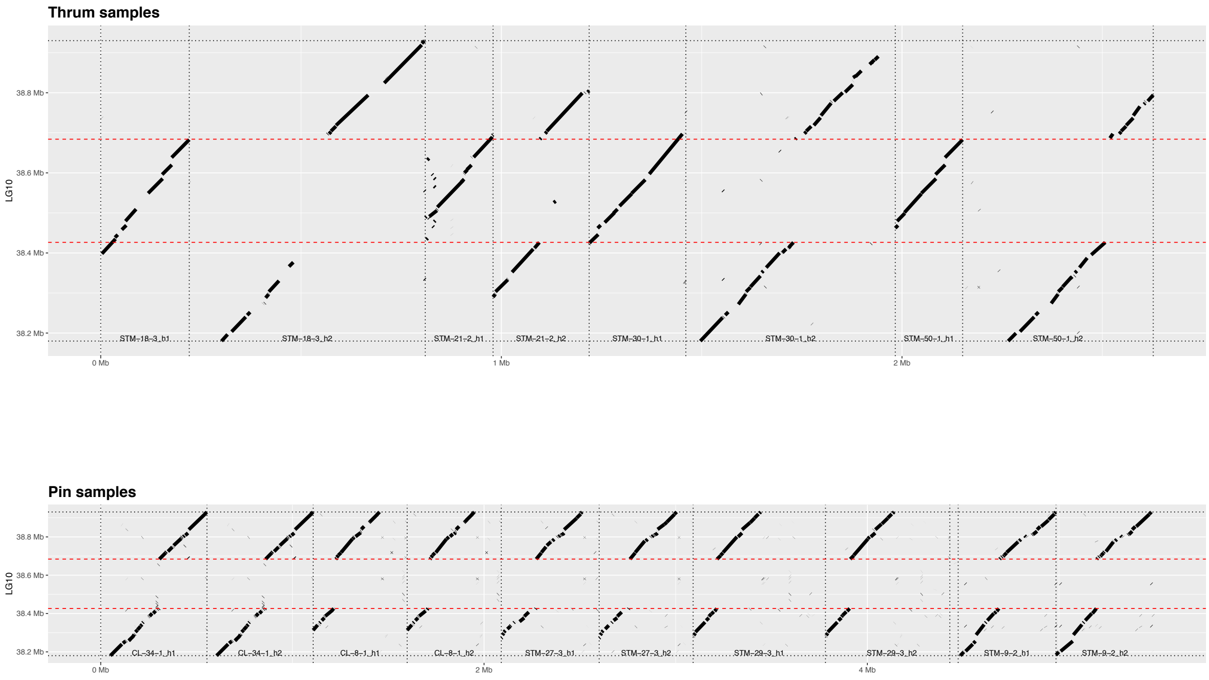


**Fig. S3. Dotplots depicting alignments between genomic assemblies for both morphs in the *S*-locus region.** Dotplots depicting the alignment between the *L. tenue* reference genome assembly region LG10: 38,180,000- 38,930,000 representing the dominant *S*-haplotype and the contigs of Supernova assemblies obtained using 10x Genomics linked-reads. Top: alignments between the candidate *S*-locus and contigs from the thrum samples (*n*=4), all of which have two haplotypes, one with and one without the thrum-specific region at the *S*-locus. Bottom: alignments between the candidate *S*-locus and contigs obtained from pin samples (*n*=5), none of which harbor the thrum-specific region. Both haplotypes are shown for each sample. Dashed red lines indicate the approximate location of the ~260 kb region that is hemizygous in thrums.

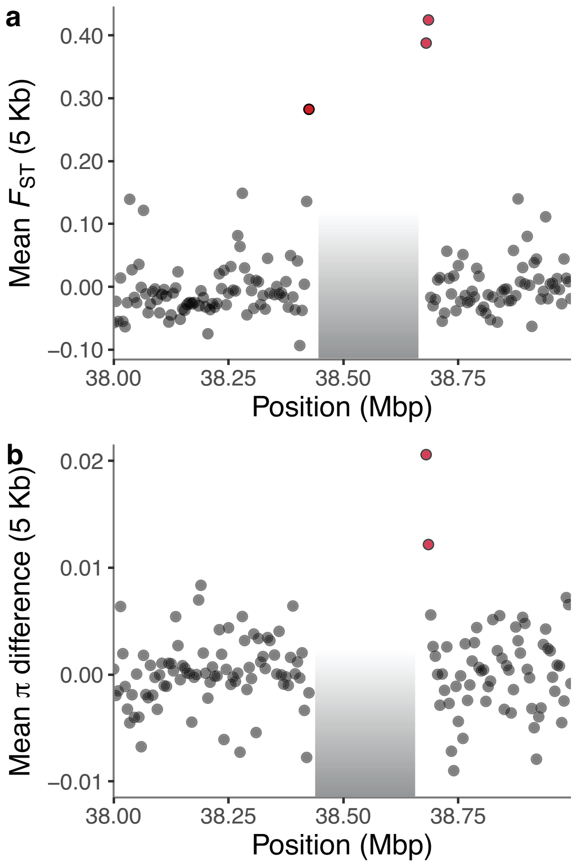


**Fig. S4. Genetic differentiation and differences in mean $\pi$ between morphs in population CL.** Estimates of **a**, genetic differentiation ($F_{ST}$) and **b**, differences in mean $\pi$ between thrum and pin sequences. **a**-**b**, Calculations were conducted on 5 kb windows across the genome (excluding the ~260 kb hemizygous region) using samples from population CL (thrum=9 and pin=8). Windows with statistically significant estimates of $F_{ST}$ and difference in $\pi$ between morphs are highlighted in red ($P < 0.01$, permutation test, 1000 replicates, followed by FDR correction with the Benjamini-Hochberg method).

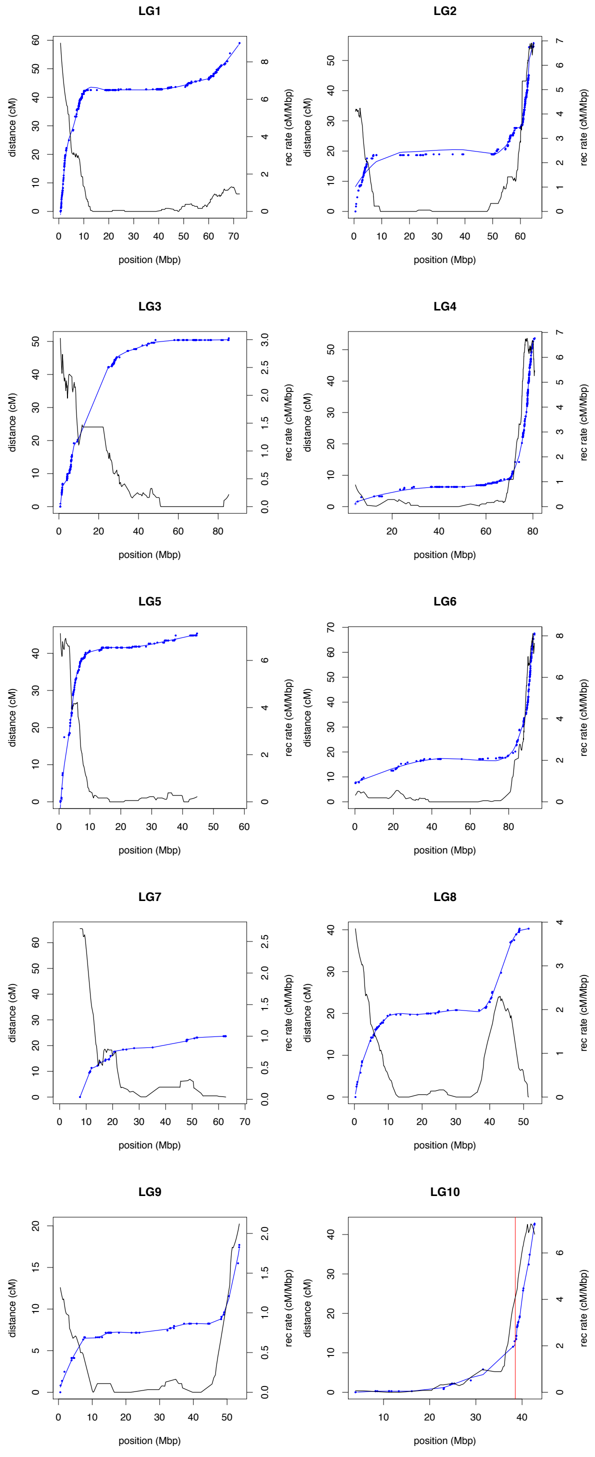


**Fig. S5. Recombination rates across LG.** Recombination rates were calculated using linkage map data for each LG. The blue dots represent genetic markers (x axis= physical position, y left axis= genetic distance in cM calculated by Lep-Map3), and the blue line depicts the local polynomial regression fitting the markers. Black lines represent recombination rates (cM/Mbp) calculated with the R package xoi (y right axis). The vertical red line in LG10 indicates the position of the the ~260 kb insertion.

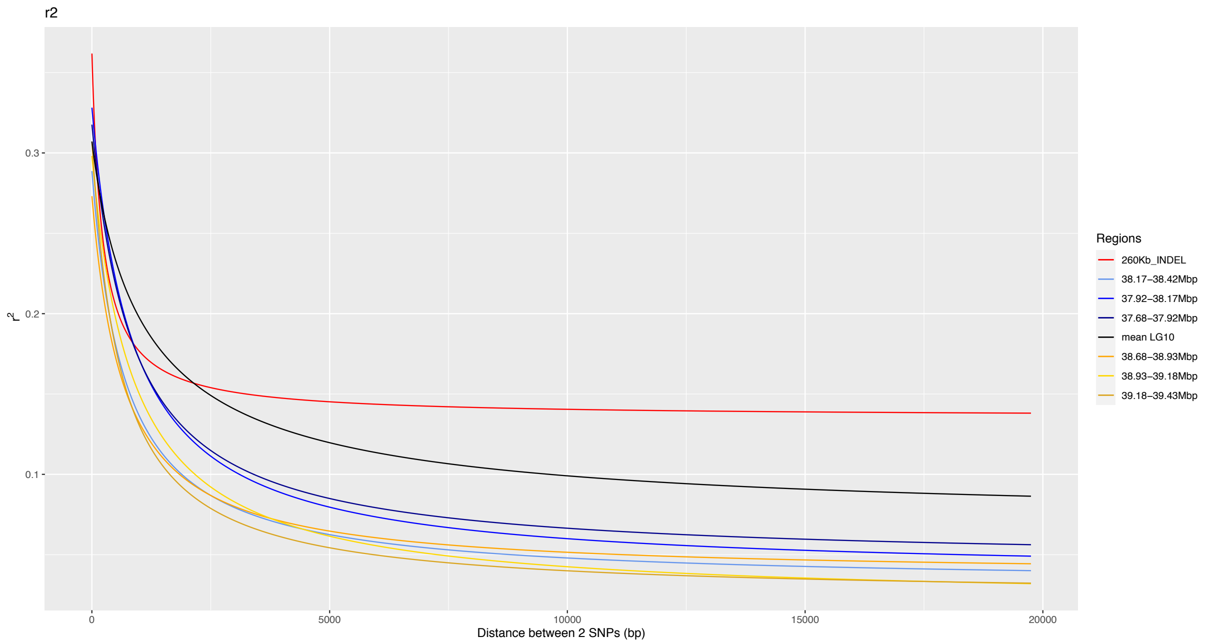


**Fig. S6. Decay of linkage disequilibrium with distance at and around the *S*-locus.** Linkage disequilibrium decay at the hemizygous region and neighboring regions. LD decay estimates were calculated with nsgLD in windows of 250 kb (*n*=43) (x axis= distance between two SNPs, y axis= $r^2$ value). LD decay is compared between the ~260 kb indel (red line), three downstream windows (blue lines= 38.17-38.42 Mbp, 37.92-38.17 Mbp, 37.68-37.92 Mbp), three upstream windows (yellow lines= 38.68-38.93 Mbp, 38.93-39.18 Mbp, 39.18-39.43 Mbp), and mean estimates across LG10-linked windows (black line).

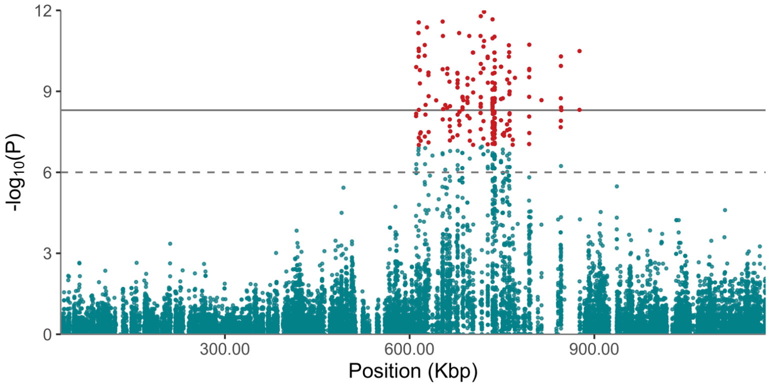


**Fig. S7. Preliminary identification of candidate *S*-locus using GWAS with the FALCON-Unzip assembly.** Association between SNPs genotypes and floral morph in *L. tenue* (CL: 8 samples of each morph, SMT: 13 samples of each morph). The Manhattan plot depicts SNPs in contig 000228Flarrow, which harbors 99.67% of loci identified as strongly associated with morph in GWAS analysis. Dashed and contiguous horizontal lines denote suggestive ($-log_{10}(1x10^{-6})$) and significant association ($-log_{10}(1x10^{-9})$) prior to *P* values correction. SNPs colored in red highlight loci that held significantly associated with floral morph following FDR correction using the Benjamini-Hochberg procedure, $P < 0.01$.

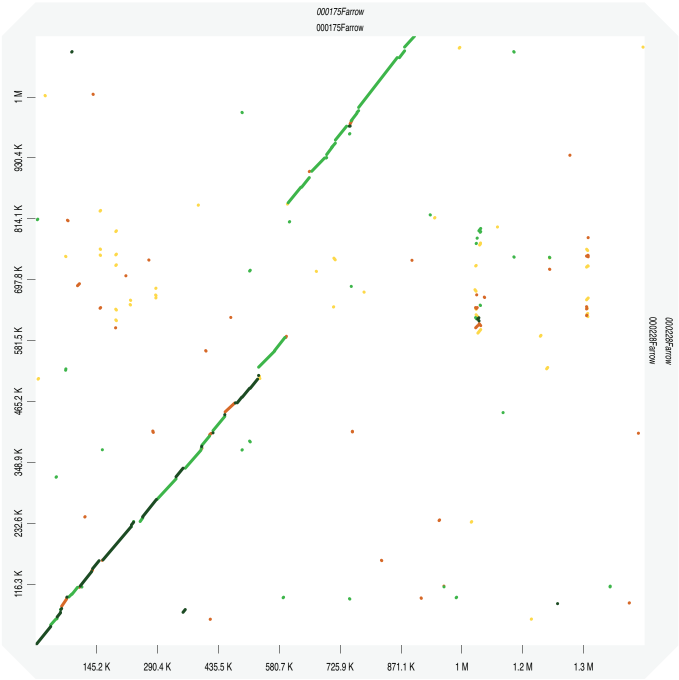


**Fig. S8. Alignment between primary contig and haplotig harboring the candidate *S*-locus.** Dot plot depicting the alignment between contigs 000175Flarrow and 000228Flarrow. Using PurgeHaplotigs, contig 000228Flarrow was identified as an haplotig of 000175Flarrow in the FALCON-Unzip version of the assembly. The dot plot was obtained with D-Genies (Cabanettes and Klopp 2018).

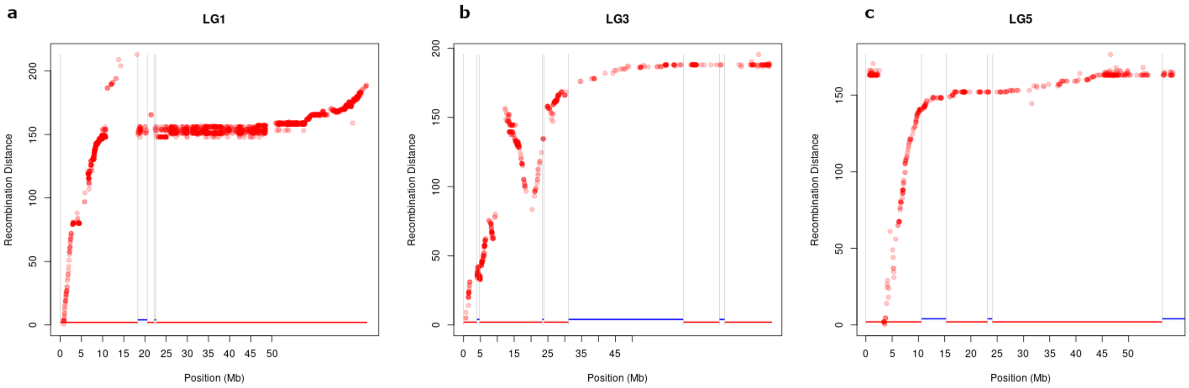


**Fig S9. Manual editing of specific LG after anchoring step.** Marey plots depicting distribution of genetic markers in LG1, LG3 and LG5 based on the inferred physical position they occupy (Mbp) and the recombination distance (cM) calculated by Lep-Map3. Blue and red lines indicate scaffolds before anchoring with Lep-Anchor.

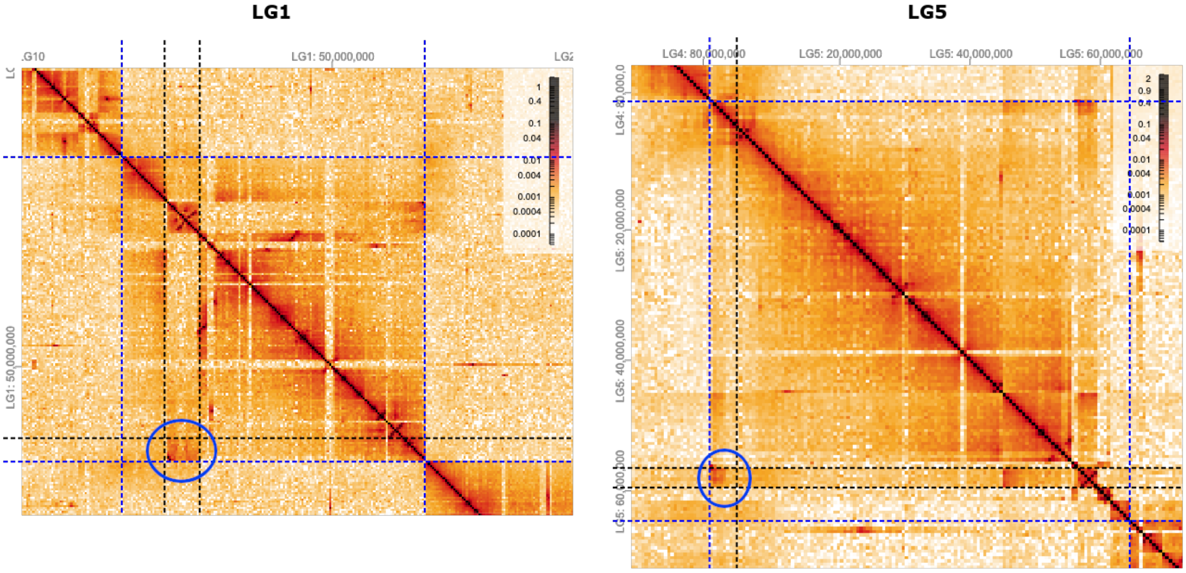


**Fig S10. HiGlass manual edit.** Heatmap of Hi-C reads mapped against the Lep-Anchor reference on LG1 and LG5. Blue dotted lines indicate the starting and ending point of linkage groups. Blue circles indicate manually edited regions: LG1:10,941,743-18,235,911 was moved to the end of LG, while LG5:1-2,929,725 was relocated to LG5:56,376,345. Color scale reflects the log-scaled number of Hi-C reads with the same barcode.

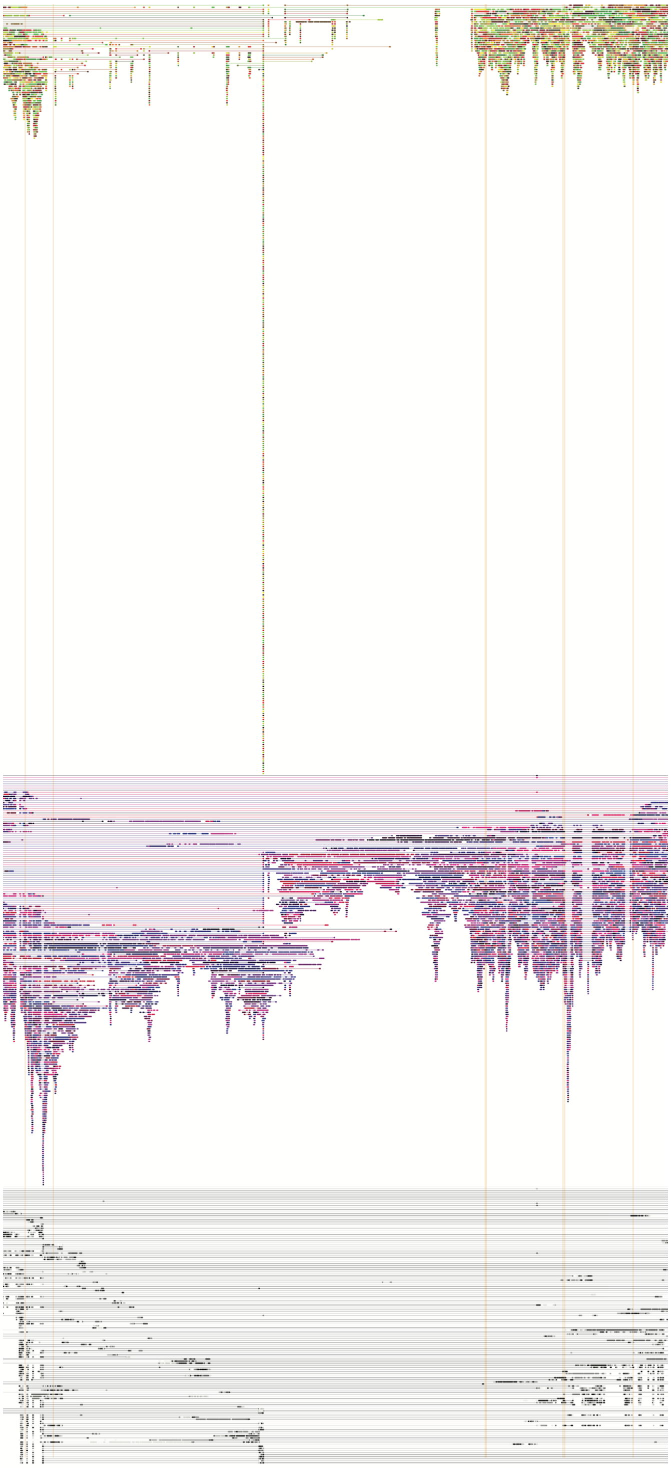

**Fig. S11. Haplotypes and indel edge determination assisted by linked-read mapping.** For all three panels, dots represent short reads, and reads aligned in the same row and in the same color correspond to the same barcode. At the top are the coordinates of LG10. Vertical orange bars denote a structural variant breakpoint called by longranger. Top: mapping of the recessive haplotype, i.e. without the ~260 kb indel. Middle: mapping of the dominant haplotype carrying with the ~260 kb indel. Bottom: unphased reads.

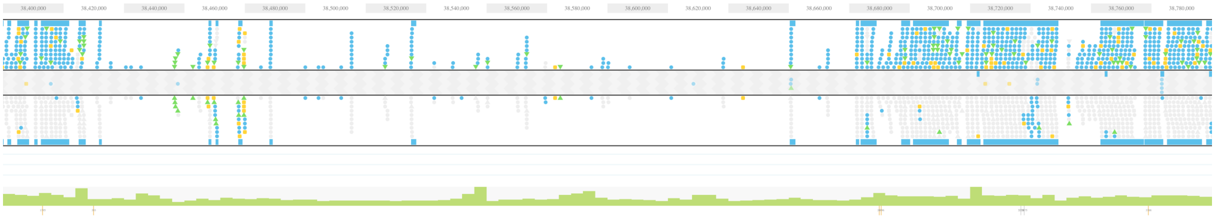

**Fig. S12. Phased haplotypes of the ~260 kb hemizygous region.** First track: recessive haplotype lacking the ~260 kb indel. Second track: dominant haplotype carrying the ~260 kb indel. Grey-colored dots represent the reference genotype while dots in other colors represent the alternative genotypes, including SNPs, small insertions and deletions. Phased heterozygous SNPs are placed at the corresponding haplotype track and unphased heterozygous SNPs are displayed in the area between the two tracks.

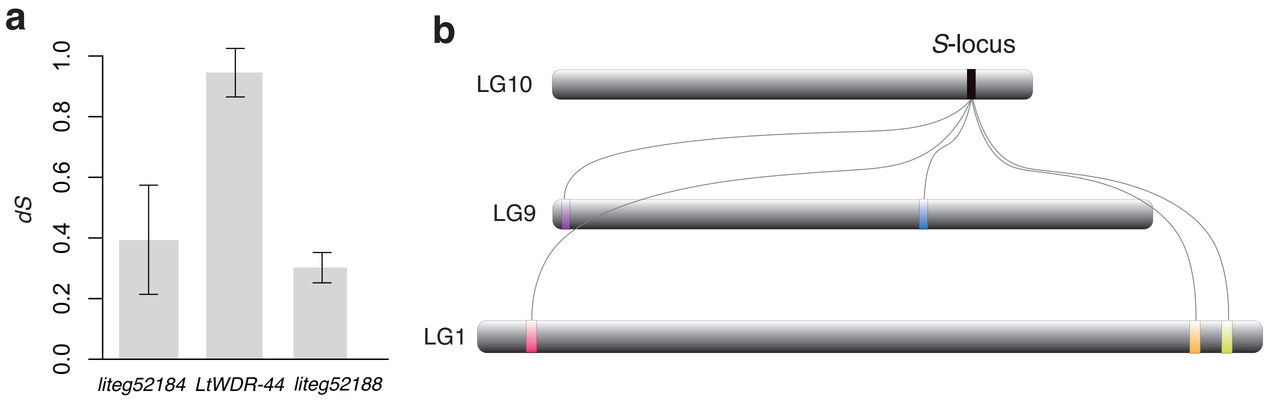


**Fig. S13. Stepwise assembly of the gene set at the *Linum* distyly *S*-locus. a.** Synonymous divergence between *S*-linked genes and their closest putative paralogs for three *S*-linked genes varies greatly, suggesting gene duplication at different times. Error bars represent standard error estimates based on 500 bootstrap replicates. **b.** Putative paralogs of *S*-linked genes are located in widely separated parts of the *L. tenue* genome assembly, suggesting stepwise assembly of the gene set at the *S*-locus.
